# Diet modulates metabolic and hepatic responses to chronic pesticide mixture exposure in mice

**DOI:** 10.64898/2026.02.18.705565

**Authors:** C. Rives, N. Poirier-Jaouen, C.M.P Martin, M. Huillet, S. Ellero-Simatos, P. Perrier, A. Polizzi, F. Lasserre, V. Alquier Bacquié, C. Guyon, Y. Lippi, C. Naylies, E.L. Jamin, N.K Dieng, R. Vuillaume, C. Orlandi, J. Gomez, S. Costes, A. Arrar, A. Lucas, S. Fried, E. Boutet-Robinet, J. Guillermet-Guibert, E. Kesse-Guyot, H. Guillou, N. Loiseau, A. Fougerat, L. Gamet-Payrastre

## Abstract

Chronic exposure to pesticide mixtures through diet is common, yet their combined metabolic effects and interactions with dietary factors remain unclear. We identified four pesticides prevalent in human exposure (imazalil, thiabendazole, boscalid, lambda-cyhalothrin) and assessed their combined impacts on hepatic metabolism and metabolic homeostasis using human liver cells and male mice fed standard chow or western diets. We found that the pesticide mixture induced metabolic perturbations in human hepatocytes. In addition, the pesticide mixture altered hepatic gene expression in chow-fed mice and exacerbated western diet-induced glucose intolerance, fasting hyperglycemia, and insulin resistance without affecting body weight or liver steatosis. These findings reveal that dietary context influences the metabolic consequences of pesticide mixtures, highlighting the need to consider nutritional status when evaluating environmental contaminant risks. Our results suggest that pesticide mixtures at reference doses may contribute to metabolic dysregulation, particularly under obesogenic dietary conditions.

**Highlights:**

- Four common pesticides in mixture disrupt metabolism in liver cells
- Dietary exposure to this pesticide mixture alters hepatic gene expression in mice
- The pesticide mixture exacerbates WD-induced disruptions in glucose homeostasis
- Pesticides and diet interact in producing the metabolic effects of a pesticide mixture

## Introduction

Extensive use of pesticides in agriculture has contributed to the contamination of air, soil, water and food, exposing humans to these substances through multiple routes. There is growing evidence that exposure to several classes of pesticides may have adverse health effects in humans, particularly metabolic disorders (Expertise Collective Inserm 2021). Epidemiological studies have established a link between occupational exposure to pesticides and a higher incidence of obesity and type 2 diabetes (T2D) (Arab and Mostafalou 2023). Dietary pesticide exposure profiles have also been associated with T2D risk in the general population (Rebouillat et al. 2022). Conversely, higher consumption of organic foods—which typically contain lower levels of pesticide residues—has been associated with reduced risks of metabolic syndrome, obesity, and T2D (Baudry et al. 2018; Kesse-Guyot et al. 2017, 2020). However, although obesity and T2D are key risk factors for metabolic liver diseases such as metabolic dysfunction–associated steatotic liver disease (MASLD), and the liver is the main organ of pollutant metabolism, the link between pesticide exposure and MASLD development is less characterized (Rajak, S. et al. 2022).

Mechanistic studies suggest that several pesticides may interfere with pathways involved in metabolic regulation and liver function (Ahmad et al. 2024; Expertise Collective Inserm 2013), notably through interactions with hepatic nuclear receptors that regulate lipid and glucose metabolism, inflammation, and detoxification processes (Capitão et al. 2017; Fujino et al. 2019; Groswald et al. 2023; He et al. 2020; Knebel et al. 2018a, 2018b; Léger et al. 2023; Lichtenstein et al. 2020; Yang et al. 2023). Moreover, several *in vitro* and *in vivo* studies have reported the pro-oxidative properties of pesticides (Jabłońska-Trypuć 2017; Rives et al. 2020; Wang et al. 2022). Consistent with this, recent data suggest that exposure to certain organophosphate pesticides is associated with biomarkers of liver injury and function in humans (Li et al. 2022). Experimental studies further indicate that individual pesticides can disrupt overall metabolic homeostasis and promote hepatic steatosis (Arciello et al. 2013; Wahlang et al. 2019; Yang and Park 2018).

Dietary exposure to multiple pesticide residues is widespread, with consumers chronically exposed to complex mixtures at levels below regulatory limits (European Food Safety Authority (EFSA) et al. 2024; Baudry et al. 2021; Castorina et al. 2003). Yet, the simultaneous presence of multiple pesticides may yield additive or more than additive effects (Wang et al. 2023), which are not predicted by single-compound assessments (Cedergreen 2014; Christen et al. 2014; de Sousa et al. 2014; Roustan et al. 2014). These complex interactions have been mostly reported in *in vitro* models (Hernández et al. 2013; Schmidt et al. 2021; Tait et al. 2022; Lichtenstein et al. 2020; Wang et al. 2023). In recent years, preclinical studies have also supported the notion that mixed pesticides can alter metabolic homeostasis and liver function (Mesnage et al. 2021); Lukowicz et al. 2018).

Despite these advances, the metabolic consequences of chronic dietary exposure to realistic pesticide mixtures, and their interaction with dietary factors remain unclear. In this study we investigated the effects of chronic dietary exposure to a relevant pesticide cocktail at toxicological references values on hepatic metabolism and energy homeostasis, and the potential interactions with dietary factors.

Based on exposure profiles identified in the NutriNet Santé cohort (Baudry et al. 2021; Rebouillat et al. 2021, 2022) and *in vitro* data, we selected four pesticides that induced, when combined, metabolic perturbations in human liver cells. The pesticide mixture was then assessed for its long-term effect in mice. The four pesticides were incorporated in a standard chow diet (CD) or in a western diet (WD) at doses allowing mice to be exposed for 20 weeks to two reference doses: the human acceptable daily intake (ADI; an estimate of the amount that can be ingested on a daily basis over a lifetime without appreciable risk to human health) or a tenfold higher dose (10ADI or 1/10 NOAEL; one-tenth of the no-observed-adverse-effect level, the highest experimentally determined dose at which no statistically or biologically significant effect has been described), for each of the four individual pesticides. Our murine findings provide evidence that exposition to toxicological reference values of each pesticide in mixture can induce molecular and phenotypic effects in a diet specific manner.

## Materials and Methods

### 1. Chemical and reagents

All pesticides used were of 98% purity (Sigma-Aldrich, France). Stock solutions at 50 mM were prepared in >99% dimethyl sulfoxide (DMSO, Sigma-Aldrich, France) and stored at −20°C. The designation ITBL 5, 15 or 30 indicates that each pesticide (imazalil [IMZ], thiabendazole [TBZ], boscalid [BSC], lambda-cyhalothrin [LCT]) in the mixture was at 5, 15 or 30 µM. LC‒MS grade methanol (MeOH) and acetic acid were purchased from Fisher Scientific (Illkirch, France). Ultrapure water was produced using a Milli-Q system (Millipore, Saint-Quentin en Yvelines, France).

### 2. Cell culture and pesticide treatments

Immortalized human hepatocytes (IHH) were a generous gift from Professor Bart Staels (Institute Pasteur, Lille, France; Samanez et al. 2012). Cells were seeded in 6- or 12-well plates precoated with 1 g/L porcine gelatin (Sigma-Aldrich) in William’s medium(Gibco) (10% decomplemented fetal calf serum, Dutscher), penicillin (100 units/mL, Sigma-Aldrich), and streptomycin (0.1 mg/mL, Sigma-Aldrich), glutamine (4 mM, Sigma-Aldrich), dexamethasone (1 nM, Supelco), and bovine insulin (8.4 nM, Sigma-Aldrich) at 37°C with 95% humidity and 5% CO_2_. Fifteen hours after seeding, cells were cultured in Dulbecco’s Modified Eagle Medium (DMEM) (pyruvate 230 µM, bovine serum albumin [BSA] 1 g/L, penicillin 100 units/mL, and streptomycin 0.1 mg/mL [Sigma-Aldrich], glutamine 4 mM [Sigma-Aldrich], dexamethasone 1 nM [Supelco], Gibco) without serum for 6 hours. Subsequently, cells were cultured in DMEM supplemented with 4% fetal calf serum, human insulin (0.1 to 1 nM) (Sigma-Aldrich), and glucose (Sigma-Aldrich) (1 mM to 4 mM) according to the experiments and treated with a 1000-fold concentrated solution of the mixture of the four pesticides (ITBL), in dimethyl sulfoxide (DMSO, Sigma-Aldrich). Controls were cells exposed to 0.1% DMSO.

### 3. Cell viability

IHH cells were seeded in 12-well plates (1 × 10^5^ cells/well) and treated either for 24 hours, 72 hours or 10 days at 5, 15, 30 µM with each pesticide alone. For the 10-day treatment, the medium was changed every 48 hours. At the end of the experiment, the cells were collected upon trypsin treatment (Sigma-Aldrich), and were counted using the Luna IITM cell counter after staining with trypan blue (Sigma-Aldrich).

### 4. Neutral lipid quantification in IHH cells and liver samples

IHH cells were seeded in 6-well plates (2.8 × 10^5^ cells/well) and treated every 2 days for 10 days with the pesticide mixture (ITBL) at 30 μM. The agonist of the liver X receptor (LXR), T-0901317 (T0, 30 μM, Sigma-Aldrich), was added 24 hours before cell recovery and was used as a positive control for intracellular neutral lipid quantification. After 10 days of treatment, cells were harvested (2 wells were pooled) by scraping in EGTA (aqueous solution) 5 mM: methanol (1:2, v/v). For mouse liver samples, the equivalent of 2 mg of tissue was homogenized in FastPrep tubes containing beads and aqueous EGTA 5 mM:methanol (1:2, v/v), then processed using a Precellys, as described previously (Bligh and Dyer, 1959).

Lipid extraction was performed by adding 2.5 mL of methanol (Sigma-Aldrich), 2 mL of Milli-Q water, and 2.5 mL of dichloromethane (Fisher Chemical) (2.5:2.5:2, v/v/v) to cell and liver samples. For cells, evaporation was performed twice before resuspending the extracts in 20 μL of ethyl acetate (Sigma-Aldrich). For liver samples, a single evaporation was performed before resuspending the extracts in 160 µL of ethyl acetate. A standard mixture composed of 6 μg of stigmasterol (2 μg/10 μL), 6 μg of cholesterol C17 (2 μg/10 μL), and 16 μg of triglycerides TG19 (4 μg/10 μL) was used. Lipids (triglycerides, free cholesterol, and cholesterol esters) were quantified by gas chromatography coupled with flame ionization detection (GC-FID) (Lipidomics Platform, I2MC, Toulouse), using a Thermo Electron system focused with a Zebron-1 Phenomenex fused silica capillary column (5 m, 0.32 mm internal diameter, 0.50 µm film thickness; Phenomenex, England), as previously described (Podechard et al. 2018). The oven temperature was programmed to increase from 200°C to 350°C at a rate of 5°C/min, and the carrier gas was hydrogen (0.5 bar). The injector and detector were set at 315°C and 345°C, respectively.

### 5. Measurement of mitochondrial oxygen consumption rate

IHH cells were treated for 24 hours with the pesticide mixture (ITBL) at 30 μM in DMEM medium (XFe24 Cell Culture Microplates pre-coated with gelatin, 6.25 × 10^5^ cells/well). Then, cells were incubated in Seahorse XF DMEM Medium, pH 7.4 (Agilent) (Seahorse XF Glucose (1 mM final), Seahorse XF Pyruvate (0.1 mM final), and Seahorse XF L-Glutamine (0.2 mM final). Real-time measurements of the oxygen consumption rate (OCR) were performed by isolating a small volume (approximately 5 µL), also known as a "transient microchamber", above the cell monolayer using the Seahorse XF Cell Mito Stress Test (Agilent, Santa Clara, CA, US).

Through an integrated drug delivery system, three compounds were sequentially added to the wells (30-minute intervals between each injection): oligomycin (2 μM, ATP synthase inhibitor), carbonyl cyanide-4 (trifluoromethoxy) phenylhydrazone (FCCP, 2 μM, mitochondrial uncoupler), rotenone/antimycin A (0.5 µM each, inhibitors of complex III of the respiratory chain), to determine ATP production, maximal respiration, and proton leak, respectively. Data were analyzed using Seahorse XFe Wave software (Agilent). The data were normalized to the cell density in each well measured by an automated IncuCyte cellular imaging system (Sartorius).

### 6. Animals and diets

*In vivo* studies were conducted in accordance with EU directive 2010/63/EU for animal experiments and approved by an independent ethics committee under authorization number 17430-2018110611093660. All mice were housed at 21–23°C with a 12h/12h light/dark cycle and had access to standard rodent chow diet (SAFE A04 U8220G10R from SAFE Augy, France) and tap water. Eight-week-old male SOPF C57BL/6 mice (Janvier Labs) were acclimatized for one week and then randomly assigned to different experimental groups. Mice were fed either a standard chow diet (CD; 70% carbohydrate, 4% fat, and 14% protein, n = 36) or a Western diet (WD; 61% carbohydrate, 20% fat, and 14% protein, n = 36) *ad libitum*. After 5 weeks, both groups were further divided into 3 subgroups: (i) a group fed a diet containing the mixture of 4 pesticides (ITBL) and exposed to the acceptable daily intake (CD-ADI or WD-ADI) of each pesticide; (ii) a group fed a diet containing the mixture of 4 pesticides (ITBL) and exposed to 10ADI, corresponding to 10 times the ADI (CD-10ADI or WD-10ADI), of each pesticide; and (iii) one group not exposed to pesticides (CD or WD). The exposure period lasted for 20 weeks (n = 12 animals/group). Rodent diets were prepared in collaboration with the SAAJ unit (Jouy-en Josas) as described previously (Lukowicz et al. 2018). The quantities of pesticides incorporated into the rodent diet were confirmed by LC-MS analysis (Eurofins, France) (supplementary Table 1). Body weight, food intake, and water consumption were monitored weekly throughout the experiment.

### 7. RNA extraction of IHH cells and liver samples

IHH cells (2 × 10^5^ cells/well, 6 well-palte) and treated for 24 hours with the mixture (ITBL) at 30 µM. The cell monolayers or the liver samples were lysed using TriReagent (MRC). After addition of chloroform (Fisher Chemical), total RNAs were extracted in the aqueous phase and then precipitated with 99.8% isopropanol (Sigma-Aldrich. After washing with 70% ethanol (Sigma-Aldrich), the RNA was resuspended in RNase/DNase-free water (Ambion). The RNA concentration was measured using a nanophotometer (Nanodrop 1000, Thermo Scientific) at an absorbance of 260 nm. The RNA was diluted to a concentration of 135 ng/µl for microarray analysis (IHH cells) or RNA sequencing (liver samples).

### 8. Gene expression analysis

#### a. Quantification of relative mRNA expression by RT-qPCR

To perform real-time quantitative PCR, 2 µg of RNA was reverse transcribed using a High-Capacity cDNA Reverse Transcription Kit (Applied Biosystems, Foster City, CA, USA). Amplification reactions were carried out in 96-well plates in a mixture consisting of SYBR Green (Low ROX SYBR MasterMix dTTP blue, Takyon), a fluorescent DNA intercalating agent, primer pairs of interest at a final concentration of 300 nM or 900 nM depending on primer efficiency, and cDNA diluted to a 1:20 ratio in ultrapure DNase/RNase-free water. The primer sequences used are presented in supplementary Table 2.

qPCR experiments were performed using an AriaMx real-time PCR system (Agilent). Fluorescence data were analyzed using LinRegPCR software to calculate the PCR efficiency per point and provide a relative initial mRNA concentration through linear regression of the exponential phase of the PCR curve. The relative expression measurements of the genes of interest were normalized to the expression level of the mRNA encoding the GAPDH protein (glyceraldehyde-3-phosphate dehydrogenase).

#### b. Gene expression profiling of IHH cells by microarray

Gene expression profiles were obtained for six different cell passages (p26 to p31) on the GeT-TRiX platform (GenoToul, Genopole Toulouse Midi-Pyrénées) using Agilent SurePrint G3 Human GE v3 DNA microarrays (8 × 60K, model 072363) following the manufacturer’s instructions. Microarray data acquisition was achieved from 200 ng of total RNA as described previously (Lukowicz et al. 2018). Microarray data and experimental details are available in the Gene Expression Omnibus (GEO) database at NCBI (GSE305353).

Microarray data were analyzed using R (https://www.R-project.org) and Bioconductor packages (Huber et al. 2015) as described previously (Lukowicz et al. 2018). Enrichment analysis for biological processes in the gene ontology (GO) was performed using Metascape (Zhou et al. 2019), and transcription factor enrichment was assessed using TRRUST.

#### c. Gene expression profiling of liver samples by RNA sequencing

For each of 66 samples, RNA-seq libraries were constructed from 1000 ng of total RNA at the GeT-TRiX facility (GénoToul, Génopole Toulouse Midi-Pyrénées) using an Illumina Stranded mRNA Prep kit (Illumina, San Diego, CA, USA) following the manufacturer’s instructions adapted to produce library sizes compatible with paired-end 150-bp read-length sequencing. The libraries were then pooled to equimolar concentrations and transferred to the GeT-PlaGe facility (GénoToul, Génopole Toulouse Midi-Pyrénées) for sequencing into one lane on an Illumina NovaSeq 6000 using a 2 × 150-bp paired-end sequencing mode with a NovaSeq 6000 S4 Reagent Kit v1.5.

Bioinformatics treatment was executed with Nextflow v23.10.0-edge (Di Tommaso et al. 2017) and processed using nf-core/rnaseq v3.14.0 (https://doi.org/10.5281/zenodo.1400710) of the nf-core collection of workflows (Ewels et al., 2020). Reads were aligned to human genome reference GRCm39 (build GCA_000001635.9, release: 2023-04). Sequencing data and experimental details are available in NCBI’s Gene Expression Omnibus (Edgar et al., 2002) and are accessible through GEO Series accession number (to be provided).

Biostatistics analyses were performed under R v4.3.0 (R Core Team, 2023) as previously described (Chousidis et al. 2025).

Clustering results are shown as a heatmap of expression signals, using the MATRIX application (Lippi and Soubès 2023), based on differentially expressed genes (p ≤0.05 and fold change >1). Gene Ontology (GO) enrichment analysis of Biological Processes was performed using Metascape (Zhou et al. 2019) with FDR <5%, using default settings.

### 9. Blood and tissue samples

Blood samples were collected from the submandibular vein into lithium heparin–coated tubes (Sarstedt, Nümbrecht, Germany) throughout the experiment (weeks 12, 16 and at the time of euthanasia). Plasma was isolated by centrifugation (1500 g, 15 min, 4°C) and stored at −80°C. Following animal euthanasia by cervical dislocation, tissue samples were collected, weighed, dissected, and used for histological analyses or frozen in liquid nitrogen and stored at −80°C until further use.

### 10. Oral glucose tolerance test (OGTT) and plasma insulin concentration

The OGTT was conducted after 23 weeks of diet. Mice were fasted for 6 hours before receiving a glucose solution (2 g/kg body weight) by gavage. Blood glucose levels were measured from the tail vein using an Accu-Check Performa glucometer (Roche Diabetes Care France, Mylan, France) 30 minutes before and 0, 15, 30, 60, 90, and 120 minutes after receiving the glucose solution. For measurements of plasma insulin concentration (see plasma biochemical analyses), 20 µL of blood was drawn from the tip of the tail vein 30 minutes before and 15 minutes after glucose gavage.

### 11. Plasma biochemical analyses

The plasma insulin concentration was measured using the We-Met platform (I2MC, Toulouse, France) with an Insulin Mouse Serum Assay HTRF kit (Revvity). During weeks 5, 12, 16, and 23, fasting blood glucose (6 hours of fasting) was measured from a drop of blood taken from the tail vein, using an Accu-Check Performa glucometer (Roche Diabetes Care France, Mylan, France). Plasma samples were analyzed to determine the levels of alanine aminotransferase (ALT), using a Cobas Mira Plus biochemical analyzer (Roche Diagnostics, Indianapolis, IN, USA) (ANEXPLO facility, Toulouse, France).

### 12. Histology

Paraformaldehyde-fixed, paraffin-embedded liver tissue sections (3 µm) were stained with hematoxylin and eosin (H&E) for histopathological analysis (n = 12 per group). The stained liver sections were analyzed blindly for steatosis. The histological features were grouped with the steatosis score evaluated according to Akpolat et al. (Akpolat et al. 2005).

Paraformaldehyde-fixed, paraffin-embedded pancreas tissue sections (4 µm–thick longitudinal sections) were stained with H&E, scanned with a Pannoramic 250 Flash III microscope, and analyzed in blinded fashion (n = 6 per group). The number of islets were quantified and the area of each islet was measured with NDP.view software, version 2.9.29 (Hamamatsu).

### 13. Analysis of pesticides and their metabolites in urinary samples

Urinary samples were collected during 24 h the week before euthanasia and stored at −80°C. Analysis of pesticides and their metabolites are described in supplementary Table 3.

### 14. Statistical analysis

Statistical analyses were performed using GraphPad Prism for Windows (version 10.2.; GraphPad Software). Data are presented as mean ± SEM. *In vitro* data were normalized to the total mean of each experiment before being pooled, with two exceptions: intracellular triglyceride measurements where the data were normalized to the number of cells in each condition before pooling; and the heatmap of genes linked to liver steatosis, hepatotoxicity, and nuclear receptor activation, where the data were normalized to control gene expression. For all experiments in IHH cells, effects were assessed with unpaired t-test, excepted for oxygen consumption rate (OCR) the effects were assessed with two-way ANOVA followed by Tuckey’s post-hoc test. For all animal experiments, differential effects were assessed with one-way ANOVA followed by Tuckey’s post-hoc test, excepted for body weight survey and the oral glucose tolerance test (OGTT) a two-way ANOVA followed by Tuckey’s post-hoc test was performed. For histology experiments and urinary metabolites analysis differential effects were assessed with Kruskal–Wallis followed by Dunn’s multiple comparisons test.

## Results

### 1. Selection of candidate pesticides

We established a list of candidate pesticides to be further evaluated for their effects on hepatic metabolism. We first used recent results from a prospective cohort that characterized the dietary exposure profiles to pesticides in a large sample of French adults with variable consumer habits. This study identified six different exposure clusters in regard to estimated dietary exposure to 25 commonly used pesticides (Rebouillat et al. 2021). In the most exposed cluster (“cluster 3”), we selected the 12 pesticides with highest estimated exposure (Rebouillat et al. 2021) (Figure 1). We next refined our selection by examining which of those 12 pesticides have a mode of action on their target organisms linked to oxidative stress and lipid metabolism (Leroux 2003; https://irac-online.org/mode-of-action/classification-online/), two key events in the development and progression of metabolic liver diseases (Friedman et al. 2018). This second selection led to a list of nine pesticides. Among them, three pesticides that were banned according to the pesticide use regulations in the European Union (EPHY – ANSES & the EU pesticide database) were excluded. The remaining six pesticides belong to four different classes: a pyrethroid insecticide (lambda-cyhalothrin), a fungicide from the strobilurin class (azoxystrobin), a fungicide from the carboxamide class (boscalid), and three azole fungicides (imazalil, thiabendazole, and tebuconazole). Finally, among the three azole pesticides, we examined those that recently showed positive association with T2D risk in the same French NutriNet-Santé cohort (Rebouillat et al. 2022), as MASLD is frequently associated with diabetes (Stefan & Cusi, Lancet Diabetes Endocrinol, 2022). This led to a final list of five pesticides, including four fungicides (azoxystrobin [AZX], boscalid [BSC], imazalil [IMZ], thiabendazole [TBZ]) and one insecticide (lambda-cyhalothrin [LCT]) (Figure 1).

**Figure 1:**
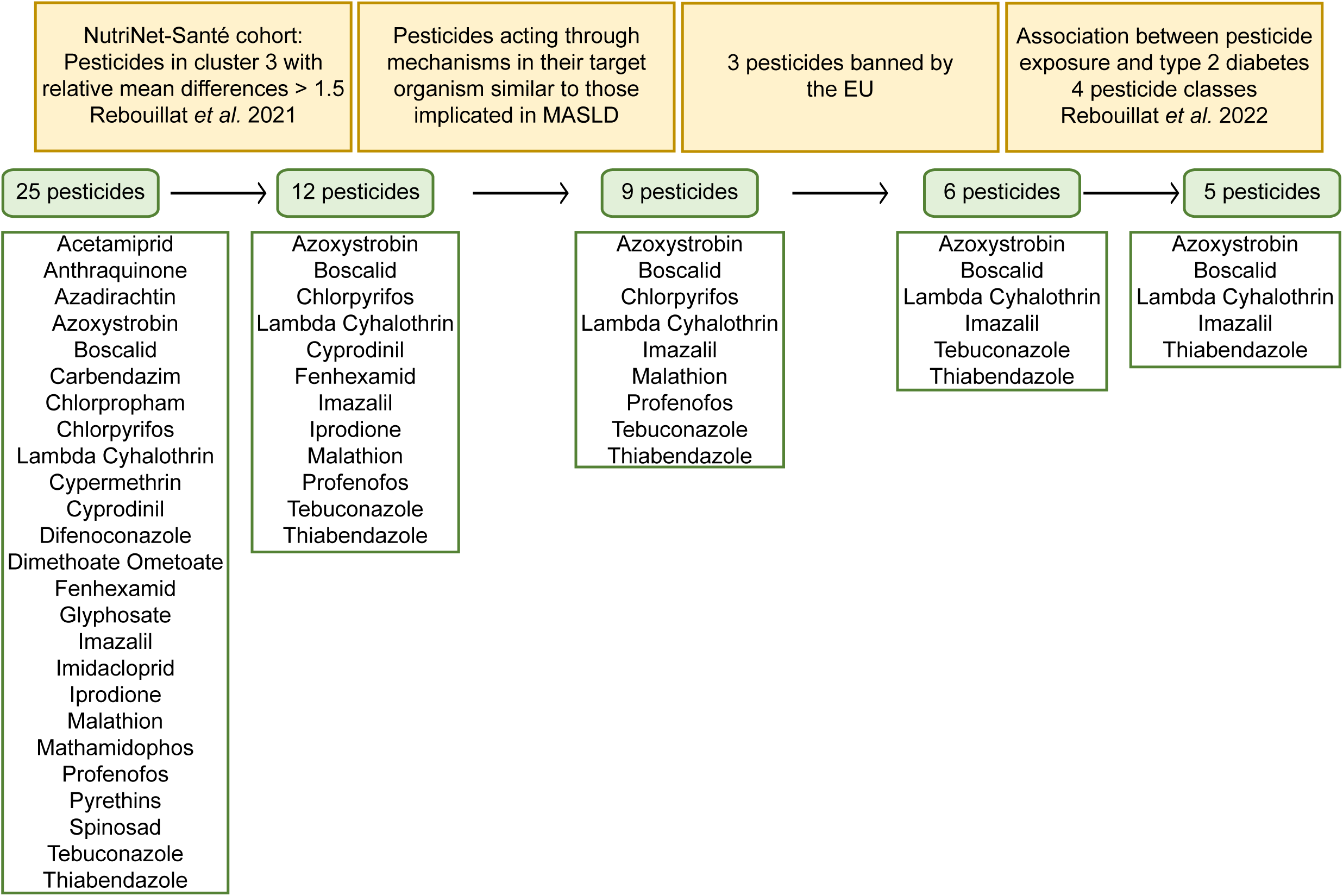
Selection of candidate pesticides.

*In vitro* exposure concentrations were derived from the ADI value of each pesticide. Estimated blood concentrations were calculated assuming a 60kg individual with a 5L blood volume, resulting in estimated concentrations in the micromolar range. Accordingly, the four compounds were tested at 5, 15 and 30 µM.

Our preliminary experiments on IHH cellular viability revealed that IMZ, TBZ, BSC, and LCT did not drastically alter cell viability in response to acute (24 h), or subchronic (72 hours or 10 days) exposure to increasing concentrations of individual pesticides (supplementary Figure 1A-C). AZX induced cytotoxicity at concentrations as low as 5 µM after 72 hours of exposure (supplementary Figure 1B) and was therefore excluded. Subsequent experiments were conducted using the four pesticides at 30 µM, the highest non-cytotoxic dose for both acute and chronic exposure in IHH cells.

### 2. Effects of the mixture of 4 pesticides on hepatic metabolism *in vitro*

We next assessed the effects of the 4 selected pesticides in mixture on IHH lipid metabolism and oxidative stress, two key events in the development and progression of obesity-associated hepatic disease (Mardinoglu 2018). We performed several *in vitro* assays in IHH cells targeting molecular initiating and key biological events of the adverse outcome pathway (AOP) for liver steatosis (AOP 34, 36, 57, 58, 517, 518 https://aopwiki.org) (supplementary Figure 2) (Mellor et al. 2016; Vinken et al. 2017).

We first evaluated the activation of the nuclear receptors PPARα, CAR, PXR, LXR and of the transcription factor AhR (the main molecular initiating event triggering liver steatosis (supplementary Figure 2)) by measuring the relative expression of their respective target genes *CYP4A11*, *CYP2B6*, *CYP3A4*, *SREBP1c*, and *CYP1A1* in IHH cells exposed for 24 h to 30 µM of each pesticide in the mixture. IHH cells exposed to the mixture of the 4 pesticides presented with a significantly higher expression of CYP2B6, CYP3A4, SREBP1 and CYP1A1 compared to untreated cells suggesting activation of CAR, PXR LXR and AhR respectively (Figure 2A).

**Figure 2:**
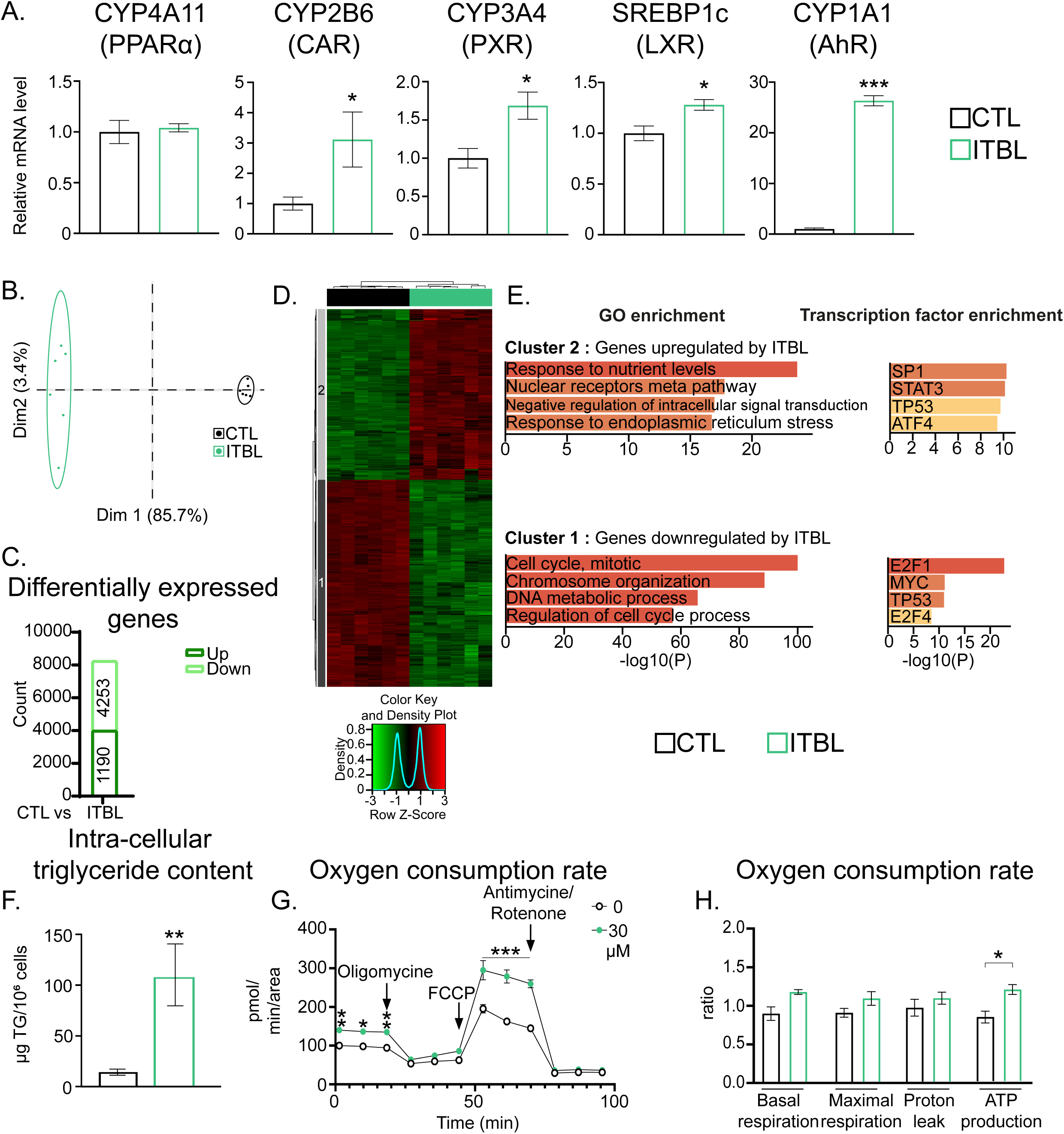
Impact of the mixture of 4 pesticides on IHH cell metabolism. **(A)** Impact of the pesticide mixture on nuclear receptor activation in IHH cells. Relative expression of CYP4A11, CYP2B6, CYP3A4, SREBP1c and CYP1A1 mRNAs in IHH cells exposed to the mixture (ITBL) at 30 µM for each pesticide for 24 h (n = 6/condition). **(B-E)** Impact of the pesticide mixture on gene expression in IHH cells. Data from a microarray experiment performed with IHH cells treated with the pesticide mixture (ITBL) at 30 µM for each pesticide in the mixture for 24 h (n = 6/condition). (**B**) Principal component analysis (PCA) score plots of the IHH transcriptomic dataset. (**C**) Number of differentially up- and downregulated genes in exposed vs. control untreated cells. (**D**) Heatmap and hierarchical clustering showing the definition of 2 gene clusters (*p* ≤ 0.05 and fold change >1.5). (**E**) Pathway and transcription factor enrichment analysis in each cluster. For each sample, the raw data were normalized to the average value of all the samples. **(F)** Impact of the pesticide mixture on triglyceride content in IHH cells. Triglyceride content in IHH cells treated with the pesticide mixture at 30 µM for 10 days (n = 3-4). **(G, H)** Impact of the pesticide mixture on mitochondrial respiration in IHH cells. (**G**) Oxygen consumption rate (OCR) profiles of IHH cells treated with the pesticide mixture at 30 µM for 24 h (n = 3-4). (**H**) Basal respiration, maximal respiration, proton leak and ATP production of IHH cells treated with the pesticide mixture. Data are presented as the mean ± SEM. * Treatment effect, * *p* < 0.05, ** *p* < 0.01, *** *p* < 0.001. (A, F, H) Unpaired parametric T test; (G)Two-way ANOVA multiple comparisons test.

Nuclear receptor activation induces changes in gene and protein expression which are considered key biological events in the steatosis AOP. Thus, we used an untargeted microarray approach to examine the whole pattern of IHH gene expression upon pesticide exposure and investigated the differences in gene expression between untreated IHH cells and those exposed to the mixture of the four compounds (Figure 2 B-E). Principal Component Analysis showed a clear discrimination between untreated and pesticide-treated IHH cells (Figure 2B). In addition, the number of differentially up- and down-regulated genes (DEGs) was increased in cells exposed to the pesticide mixture compared to control non-exposed cells (Figure 2C). Hierarchical clustering of DEGs (p<0.05 and fold change >1.5, 4205 genes) highlighted two clusters with gene expression levels that differed between unexposed IHH cells and those exposed to the pesticide mixture (Figure 2D). Genes from cluster 1 were downregulated in cells exposed to the pesticide mixture and were linked to cell cycle processes and enriched in E2F1/4, MYC, and TP53 target genes. Genes from cluster 2 were upregulated in cells exposed to the pesticide mixture and are mainly targets of SP1, STAT3, TP53, and ATF4 transcription factors. The top related biological functions were “response to nutrient levels”, “nuclear receptor meta pathway”, “negative regulation of intracellular signal transduction”, and “response to endoplasmic reticulum stress” (Figure 2E). We then focused our analysis on relevant genes involved in liver steatosis, nuclear receptor activation, and hepatotoxicity (Lichtenstein et al. 2020). As shown in supplementary Figure 3, IHH exposure to the pesticide mixture led to significant upregulation of a large number of these genes, relative to gene expression in untreated cells. The most strongly induced genes are involved in xenobiotic metabolism (*SULT1C2*, *CYP3A5*, *UGT2B7*, *CYP2B6*, *POR*) and in lipid metabolism (*PNPLA3*, *SCD1*, *SREBF1*, *MSMO1*, *PPARa*, *HADHB*), supporting the potential pro-steatotic impact of the pesticide mixture.

We next measured intracellular triglyceride levels in IHH cells treated for 10 days with the pesticide mixture using GC-FID as lipid accumulation is a key event in the AOP for liver steatosis and a hallmark of the disease. Exposure of IHH cells to the pesticide mixture at 30 µM resulted in a higher content of triglycerides compared with that of untreated cells (Figure 2F).

At the organelle level, mitochondrial disruption has been proposed to be a late key event in the steatosis AOP. Thus, we next evaluated the effect of combined pesticides on IHH cell mitochondrial respiratory functions using Seahorse XF stress test technology. Exposure of IHH cells to the pesticide mixture led to an increased basal and maximal mitochondrial respiration (Figure 2G) and a significant rise in ATP production (Figure 2H).

Altogether, combining data from previously published epidemiological studies and a panel of *in vitro* assays, we identified four commonly used pesticides that, when combined, induced metabolic perturbations in liver cells.

### 3. Chronic dietary exposure to the mixture of 4 pesticides in mice

We next investigated the *in vivo* metabolic and hepatic effects of the pesticide mixture and the interactions with dietary factors. Adult male mice were first fed either a control diet (CD) or a western diet (WD) for 5 weeks and then exposed to the pesticide mixture through these diets for an additional 20 weeks (Figure 3A). Pesticides were incorporated in the CD and WD at doses exposing mice to the ADI (CD- or WD-ADI) or 10 times ADI (CD- or WD-10ADI) of each of the four pesticides in the mixture (Figure 3A). Pesticide levels quantified in the feed pellets confirmed that the concentration of the four pesticides in each diet was close to the expected quantities (supplementary Table 1).

**Figure 3:**
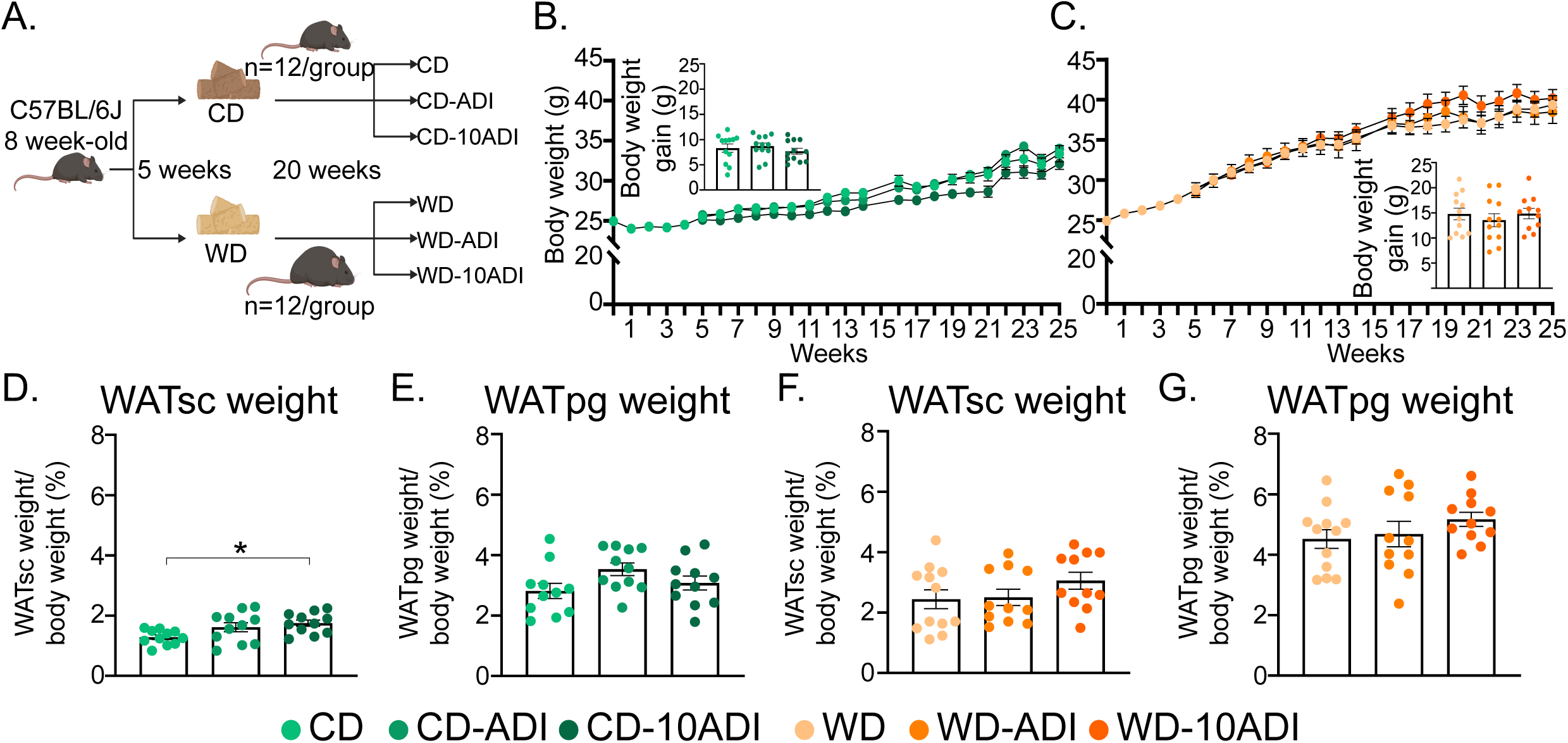
Pesticide mixture exposure does not influence mouse body weight, regardless of diet. **(A)** Experimental design. Eight-week-old male C57BL6J mice were fed a control diet (CD) or a Western diet (WD) for 5 weeks. Both groups were then divided into 3 subgroups: one fed a diet containing the mixture of 4 pesticides (ITBL) and exposed to the acceptable daily intake (CD- or WD-ADI) of each pesticide; one fed a diet containing the mixture of 4 pesticides (ITBL) and exposed to 10 times the ADI (CD- or WD-10-ADI) of each pesticide; and one not exposed to pesticides (CD or WD) for 20 weeks (n = 12 animals per group). **(B, C)** Body weight in each group from week 0 prior to exposure through 25 weeks. The bar graphs show the body weight gain at the end of the experiment. **(D-G)** Relative subcutaneous (sc) (**D**, **F**) and epididymal (pg) (**E**, **G**) white adipose tissue weight in each group of mice (n = 12 per group). Data are presented as the mean ± SEM (body weight follow-up (B, C), two-way ANOVA followed by a Tuckey’s post-hoc test; bar graphs (B-G), one-way ANOVA followed by a Tuckey’s post-hoc test).

Body weight did not show any significant differences between exposed (ADI and 10ADI) and non-exposed mice in both the CD- and the WD-fed groups (Figure 3B, C). Perigonadal (WATpg) and subcutaneous (WATsc) white adipose tissue weights were also not significantly changed by pesticide mixture exposure in mice fed a CD or WD, except for a small increase in the relative WATsc weight in the CD-10ADI compared with that in the CD mice (Figure 3D-G). Food and water intake also did not differ between exposed and unexposed mice in both the CD- and the WD-fed mice (supplementary Figure 4A, B).

We next evaluated the impact of pesticide mixture exposure on liver homeostasis during CD and WD feeding. Animal exposure to the pesticide mixture did not impact liver weight nor induce hepatic damage, whatever the dose and the type of diet (CD or WD) (Figure 4A-D). Histological analysis of H&E-stained liver slices and hepatic triglyceride quantification confirmed the absence of pesticide impact on hepatic steatosis in WD-fed mice (Figure 4E, F). By contrast, a slight but significant increase in the steatosis score and a trend toward higher hepatic triglyceride levels were observed in CD-ADI–fed mice compared with unexposed mice (Figure 4E, F).

**Figure 4:**
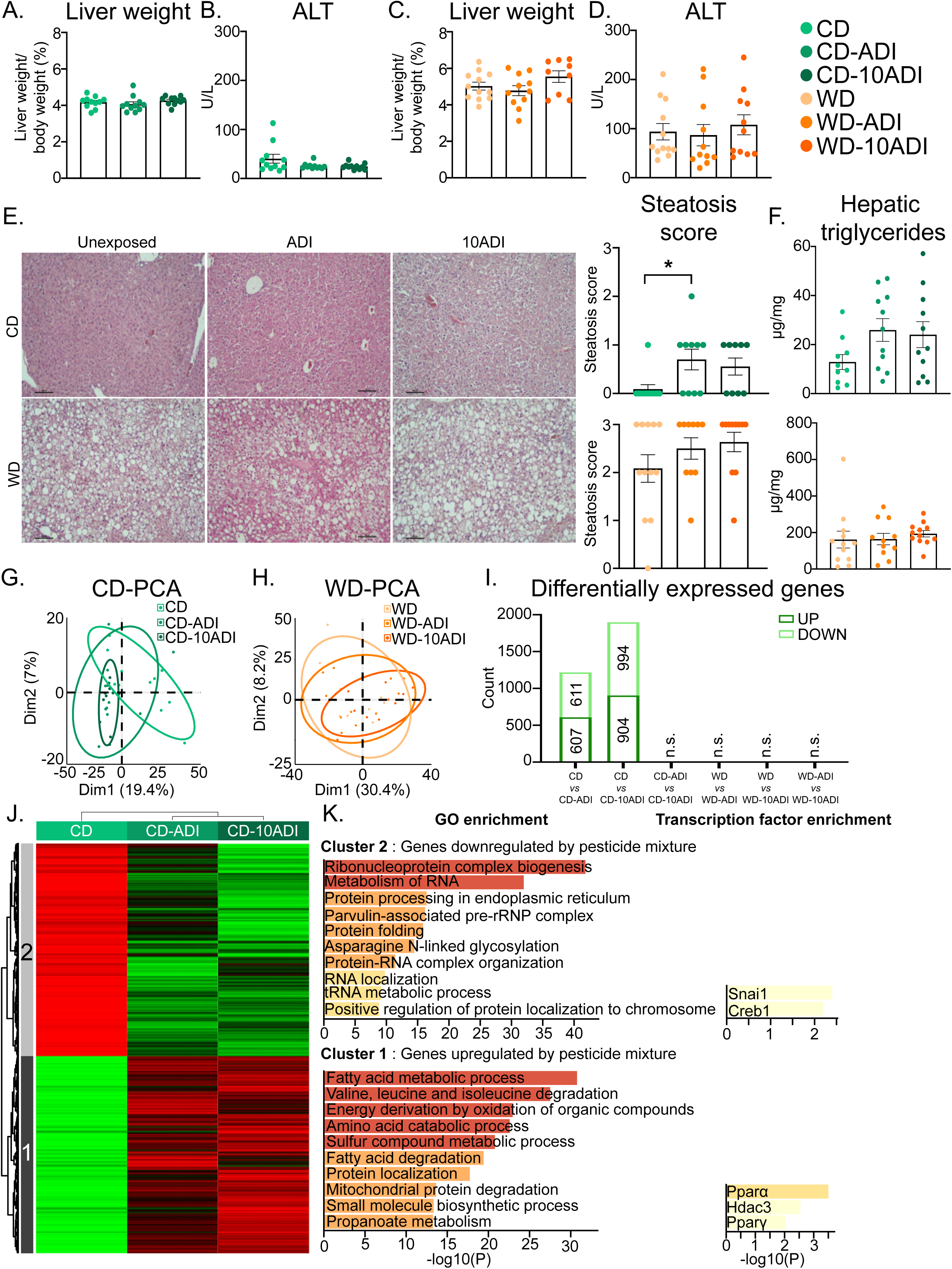
Pesticide mixture exposure changes hepatic gene expression in a diet-dependent manner. **(A-D)** Relative liver weight and plasma alanine aminotransferase (ALT) levels of CD-fed (**A**, **B**) and WD-fed (**C**, **D**) mice in each group (*n* = 12 per group). **(E)** Representative histological sections (magnification ×100) of liver stained with hematoxylin and eosin (H&E) and estimated liver steatosis score in each group (*n* = 12 per group). **(F)** Hepatic triglyceride content in each group (*n* = 12 per group). **(G-K)** Data from an RNA-seq experiment performed with liver samples from each group of mice (*n* = 8/group). Principal component analysis (PCA) score plots of liver transcriptomic dataset in CD- (**G**) and WD-fed mice (**H**). Number of differentially up- and downregulated genes in CD- and WD-fed mice unexposed vs. exposed to ADI, unexposed vs. exposed to 10ADI or exposed to ADI vs. exposed to 10ADI (n.s., nonsignificant) (**I**). Hierarchical clustering showing the definition of 2 gene clusters (*p* ≤ 0.05 and fold change >1) (**J**) and pathway and transcription factor enrichment analysis in each cluster (**K**). Data are presented as the mean ± SEM. * Treatment effect, * *p* < 0.05, one-way ANOVA followed by Tukey’s post-hoc test (A-D, G); Kruskal-Wallis followed by Dunn’s post-hoc test (F).

To further explore the potential impact of pesticide exposure on the liver, we analyzed the hepatic transcriptome in each animal group using RNA sequencing. PCA of gene expression profiles showed a slight separation between exposed and non-exposed mice fed a CD along the first principal component, accounting for 19.4% of the variance (Figure 4G). By contrast, PCA did not allow discrimination between exposed and non-exposed WD-fed mice, whatever the dose of pesticide mixture (Figure 4H). The number of differentially up- and down-regulated genes in CD-fed mice exposed to the ADI of each pesticide in the mixture (607 up- and 611 down-regulated genes) and to 10ADI of each pesticide in mixture (904 up- and 994 down-regulated genes) was higher than it was in unexposed animals (Figure 4I). By contrast, exposure to the pesticide mixture in WD-fed mice did not affect the number of DEGs compared with that in the unexposed animals. We performed hierarchical clustering of DEGs (p<0.05 and fold change >1; 2311 genes) in the CD, CD-ADI, and CD-10ADI groups (Figure 4J). Two clusters of genes were identified. Genes from clusters 1 and 2 were respectively up-and down-regulated in exposed CD-fed (CD-ADI and CD-10ADI) compared with their expression in unexposed CD-fed mice (Figure 4J). Upregulated genes are linked to fatty acid metabolism, amino acid metabolism, and cellular respiration, and are mainly enriched in PPARα targets. Downregulated genes from cluster 2 are mainly involved in RNA and protein processing (Figure 4K).

Altogether, liver analysis showed that exposure to the pesticide mixture did not exacerbate WD-induced alterations in hepatic phenotype and gene expression. However, exposure to the pesticide mixture in CD-fed mice was associated with significant changes in hepatic gene expression.

To further investigate the differential hepatic impact of pesticide exposure according to the type of diet, we compared pesticide metabolism in CD- and WD-fed mice. Analyses of urine samples by UHPLC-HRMS allowed the detection of several pesticides and their metabolites. As shown in Figure 5A-D and supplementary Table 3, TBZ and BSC and 3 of their metabolites were detected, including phase 1 metabolites (boscalid 5-hydroxy, thiabendazole 5-hydroxy) and phase 2 metabolites (glucuronide and sulfate conjugates). As expected, all pesticide metabolites were found at higher levels in urine of animals exposed at 10 times the ADI than in urine from animals exposed to the ADI. Overall, more pesticide metabolites or higher levels were detected in exposed CD-fed mice than in exposed WD-fed mice (figure 5A-D).

**Figure 5:**
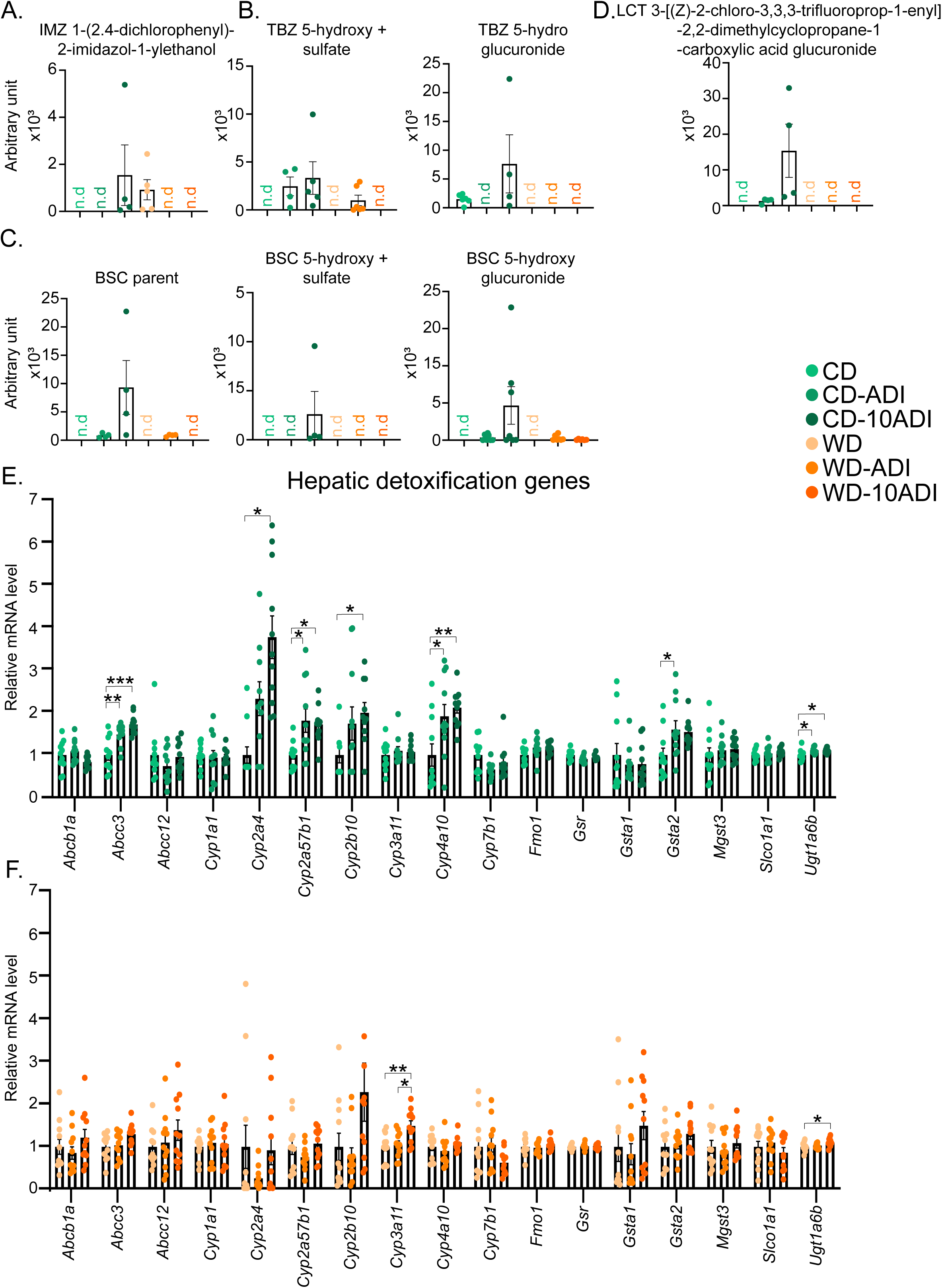
Urinary pesticide metabolites and mRNA expression level of genes involved in xenobiotic metabolism. **(A-D)** Normalized intensities of one metabolite of imazalil (**A**), two metabolites of thiabendazole (**B**), boscalid and two of its metabolites (**C**), one metabolite of lambda-cyhalothrin (**D**) measured by UHPLC-HRMS in 24 h urine samples of each group of mice (n = 8 per group). Data are presented as the mean ± SEM. *Treatment effect, \**p* < 0.05, \*\**p* < 0.01 (Kruskal-Wallis followed by Dunn’s post-hoc test); n.d.: non-detected metabolites. **(E, F)** mRNA expression of hepatic genes involved in xenobiotic metabolism in CD-(**E**) and WD-fed (**F**) mice exposed and non-exposed to the pesticide mixture (n = 8 per group). Data are presented as the mean ± SEM. *Exposed vs. non-exposed mice, \**p* < 0.05, \*\**p* < 0.01 (one-way ANOVA followed by Tuckey’s post-hoc test).

To evaluate the pesticide-detoxifying capacities of CD- and WD-fed mice, we then focused the hepatic RNA sequencing analysis on genes involved in xenobiotic metabolism. The expression profiles of hepatic genes encoding xenobiotic metabolism enzymes in the six experimental animal groups are presented in Figure 5E, F. They reveal that exposure to the pesticide mixture significantly increased the expression of 7 genes in mice fed a CD compared with only 2 in mice fed a WD.

Taken together, these results suggest differences in the pharmacokinetics of pesticides, especially in metabolism, between exposed CD- and WD-fed mice. To determine whether extrahepatic pesticide metabolism occurs in WD-fed mice—for example in adipose tissue, which can store lipophilic pollutants—we analyzed WAT for the expression of genes encoding enzymes that regulate detoxification, and of other genes involved in lipolysis, adipogenesis, glucose metabolism, and inflammation (supplementary Figure 5). None of these genes’ expression was significantly impacted by pesticide exposure in CD- or WD-fed mice. Similarly, brown adipose tissue (BAT) gene expression of BAT markers and batokines did not significantly differ between pesticide-exposed and unexposed mice, both under WD and CD (supplementary Figure 6). We also evaluated the impact of pesticide mixture exposure on the digestive tract as the first target of dietary pollutants. Expression analysis of genes involved in the structural integrity and permeability of the ileum of the intestine revealed that males fed a CD and exposed to 10 times the ADI of pesticides in a mixture had reduced expression of several genes involved in the ER stress response (*Xbp1s*), antimicrobial activity (*Reg3b* and *Reg3g*), and permeability (*Cldn2*) (supplementary Figure 7). However, the expression of none of these genes was significantly affected by pesticide exposure in WD-fed mice. Together, these results indicate diet-dependent differences in pesticide metabolism, suggesting that the bioavailability of pesticides and/or the animals’ detoxifying capacity differ according to the nutritional context.

We next evaluated the consequences of chronic exposure to the pesticide mixture on glucose homeostasis. At week 18 of exposure, glucose tolerance, fasting glycemia and insulinemia, and HOMA-IR were not affected by pesticide mixture exposure in mice fed a CD (Figure 6A-D). In contrast, WD-fed mice exposed to the pesticide cocktail at 10ADI exhibited significantly higher glucose intolerance, fasting glycemia, insulinemia, and HOMA-IR compared with animals fed the WD but unexposed (Figure 6E-H). To further investigate pesticide mixture–induced glucose homeostasis perturbations in WD-fed mice, we analyzed pancreatic endocrine mass and islet number. Although both parameters were significantly higher in the WD-fed mice than in the CD-fed mice, they did not differ significantly between exposed and unexposed animals under WD (Figure 6I, J). Overall, these results show that dietary exposure to the mixture of pesticides amplified WD-induced glucose homeostasis perturbations that were not associated with a compensatory increase in pancreatic endocrine mass.

**Figure 6:**
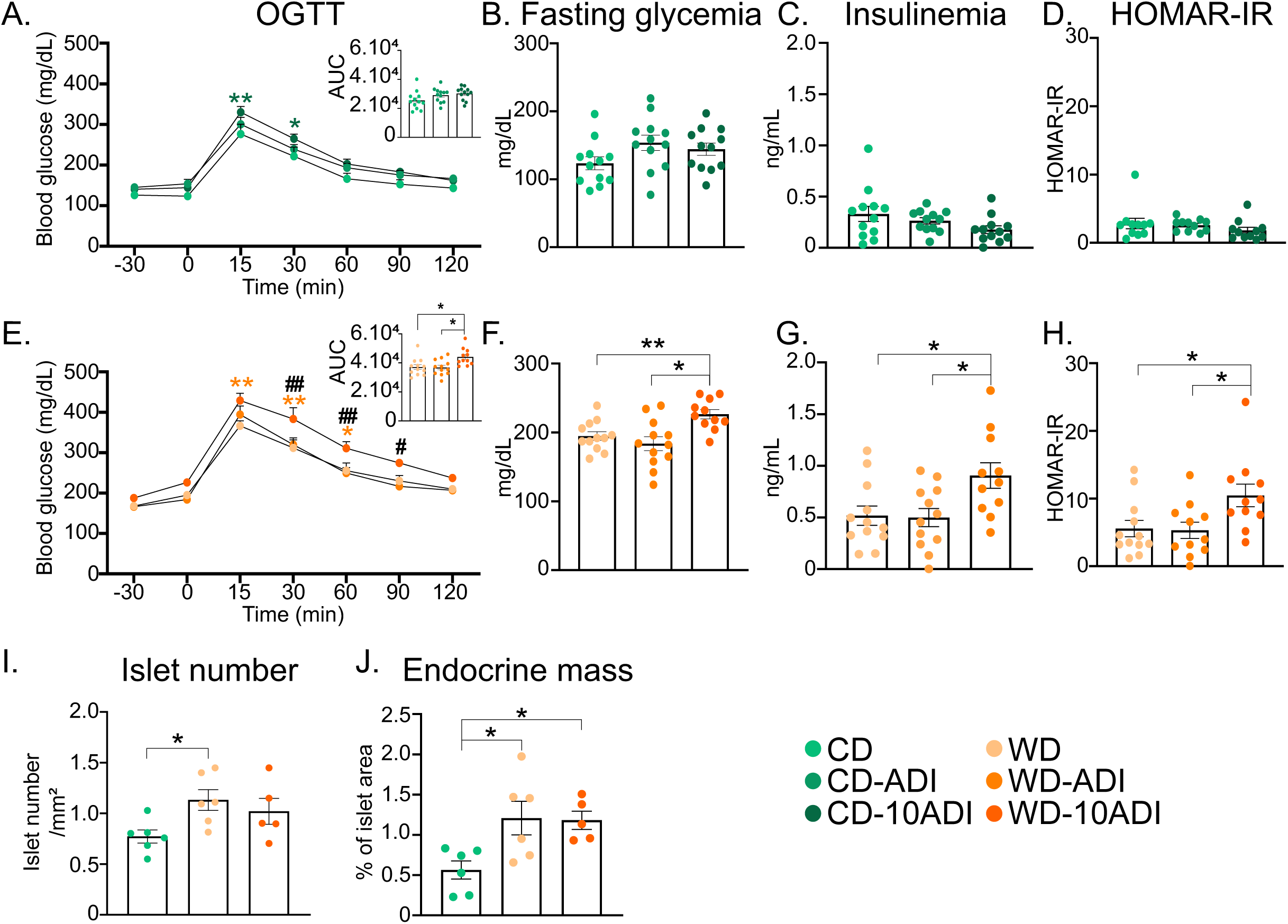
Pesticide mixture exposure exacerbates WD-induced glucose homeostasis perturbations. **(A, E)** Oral glucose tolerance test (OGTT) performed after 18 weeks of pesticide exposure in CD- (**A**) and WD-fed (**E**) mice in each group (*n* = 12 per group) and area under the curve (AUC) representing OGTT results. **(B, F)** Fasting glycemia. **(C, G)** Fasting insulinemia. **(D, H)** HOMAR-IR. **(I)** Islet number per mm^2^ of pancreas. **(J)** Percent of section area occupied by islets. Data are presented as the mean ± SEM. * Exposed vs. non-exposed mice, * *p* < 0.05, ** *p* < 0.01; # exposed ADI vs. exposed 10ADI, # *p* < 0.05, ## *p* < 0.01 (Two-way ANOVA followed by Tukey’s post-hoc test (A, E); One-way Anova followed by a Tuckey’s post-hoc test (bar graphs A-J).

## Discussion

In this work, we aimed to evaluate the metabolic consequences of a realistic dietary exposure to a mixture of pesticides at their regulatory reference doses, and to determine whether these effects vary depending on the diet composition. The innovative aspects of our study lies in (i) the integration of human epidemiological exposure profiles to guide our pesticide selection (Baudry et al. 2021; Rebouillat et al. 2021), (ii) the use of several *in vitro* assays to identify a pesticide mixture that impact metabolic processes in liver cells, (iii) an *in vivo* study assessing the pertinence of the toxicological reference values (the ADI and 10ADI; corresponding to 1/10 NOAEL) of the 4 selected pesticides in mixture and the influence of diet composition on pesticide mixture-induced effects.

The aim of our *in vitro* strategy was to characterize the metabolic effects of the pesticide mixture in liver cells. Our main findings were that the selected pesticide mixture induced gene expression changes, triglyceride accumulation and mitochondrial activity perturbations in IHH cells suggesting its pro-steatotic potency.

Our *in vivo* study revealed that the metabolic effects of the pesticide mixture in mice were not uniform, but rather dependent on the type of diet (CD vs WD), highlighting the role of nutritional context in shaping toxicological outcomes. While dietary exposure to the pesticide mixture did not elicit significant alterations in the body weight of mice fed either a CD or a WD, it led to a slight but significant increase in the steatosis score in CD-fed mice. The nonsignificant changes in hepatic triglyceride levels in CD fed mice exposed to the pesticide mixture is not entirely consistent with our *in vitro* studies, which showed a significant increase in triglyceride content in IHH cells upon exposure to the pesticide mixture. However, the changes observed in the hepatic gene expression profile of mice fed a CD support a pro steatotic property of the pesticide mixture. Pathway enrichment analysis of the hepatic transcriptome of CD-fed mice identified fatty acid metabolic processes as the top biological function associated with up-regulated genes in response to pesticide mixture exposure. It cannot be excluded that the 20-week duration of pesticide exposure in our *in vivo* experiment was insufficient to induce detectable phenotypic alterations. In our previous study, we demonstrated that pesticides induced steatosis after 6 months of exposure (Lukowicz et al. 2018). This observation is consistent with other findings that reported pesticide-induced liver damage in mice fed a control diet over extended periods (Dinca et al. 2023; Docea et al. 2018, 2019; Fountoucidou et al. 2019). The discrepancy between *in vivo* and *in vitro* results may stem from interspecies differences or/and from the inability of *in vitro* models to fully capture the complex whole-organism liver responses shaped by interorgan interactions, including those with adipose tissue and the gut microbiota (Djekkoun et al. 2021; Nichols et al. 2024; Velmurugan et al. 2017; Wang et al. 2021).

While dietary exposure to the pesticide mixture of WD-fed mice did not induce liver phenotypic and genomic alterations, it exacerbated WD-induced diabetic symptoms, including fasting hyperglycemia, glucose intolerance, and insulin resistance at tenfold ADI dose. These data suggest that exposure to this pesticide mixture may contribute to the development of glucose metabolism disruption in the presence of other risk factors, such as chronic consumption of high-fat foods. Our findings agree with recent studies showing that high-fat dietary intake may enhance the metabolic effects of pesticide exposure. Oral administration of cypermethrin to adult male mice disrupts glucose homeostasis and induces prediabetic symptoms in high-fat diet-fed animals (Wei et al. 2023a). Other studies in rodents have demonstrated that the metabolic perturbations induced by a high-fat diet are potentiated by exposure to low doses of permethrin, chlorpyrifos, perfluorooctanoic acid, and bisphenol A (Attema et al. 2022; Li et al. 2023; Ma et al. 2021; Wang et al. 2021; Xiao et al. 2018). The interaction between dietary factors and pesticide exposure has primarily been demonstrated for individual compounds, with effects varying based on the specific contaminant and exposure duration. However, studies investigating the interplay between diet and pesticide mixtures remain scarce. In a zebrafish model, exposure to a mixture of organochlorine pesticides exacerbated the diabetogenic consequences of a high-fat diet (Lee et al. 2023). Our study extends these findings in mammals, showing that a pesticide mixture can exacerbate WD-induced disruption of glucose homeostasis.

Unlike their expression levels in CD-fed mice, the expression levels of xenobiotic metabolizing enzymes were not increased in livers of WD-fed mice following exposure to the pesticide mixture, suggesting reduced pesticide metabolism or the occurrence of extrahepatic pesticide metabolism in WD-fed mice. As urinary profiles of pesticide metabolites were similar between non-exposed and pesticide-exposed WD-fed mice, we hypothesize that pesticides are overall less metabolized in WD-fed mice and may accumulate in other tissues. Previous studies reported that several pesticides, because of their lipophilicity, target adipose tissue (Barrios-Rodríguez et al. 2021; Chang et al. 2016; Jackson et al. 2017; Sousa et al. 2023). Complementary experiments would be necessary to fully elucidate the fate of pesticides in WD-fed mice.

The observed hyperglycemia and insulin resistance in WD-fed mice exposed to the pesticide mixture may not be attributed to alterations in gluconeogenesis or glycogen synthesis or decreased glucose uptake in insulin-sensitive tissues, as observed in other studies (Wei et al. 2023b). Indeed, while we did not directly measure hepatic glucose uptake, we found that exposure to the pesticide mixture did not affect the expression of genes involved in glucose synthesis and transport or in insulin signaling in the liver (results not shown). This suggests that the liver may not be the primary target tissue through which the pesticide mixture influences glucose metabolism in WD-fed mice. Whether glucose uptake by skeletal muscle and/or adipose tissue is impaired upon pesticide mixture exposure remains to be determined.

T2D occurs when β cells fail to adequately increase insulin secretion to meet demands to counteract insulin resistance, and this failure may be exacerbated by a reduction in β-cell mass over time (Costes et al. 2021). Several pieces of evidence indicated that pancreatic β cells may be targeted by pollutants (Hectors et al. 2011; Hoyeck et al. 2022; Lee et al. 2017). In this study, fasting hyperglycemia and increased fasting insulin in response to WD were both higher in mice exposed to pesticide (10ADI or 1/10 NOAEL) than in non-exposed mice, suggesting that the increased secretory function of pancreatic islet β cells was not sufficient to compensate for insulin resistance. In addition, WD-fed mice exposed to the pesticide mixture showed no differences in islet number and endocrine mass when compared with unexposed mice. While we cannot exclude a difference between the numbers of alpha and beta cells, these data suggest that in WD-fed mice exposed to the pesticide mixture, the endocrine pancreas fails to counteract pesticide-induced glucose intolerance and insulin resistance, in contrast to what occurred in males fed a WD but not exposed.

In conclusion, our study demonstrates that chronic dietary exposure to reference doses of each pesticide of this realistic cocktail is associated with significant changes in hepatic gene expression in CD-fed mice and exacerbates WD-induced disruption of glucose homeostasis. This is one of the few studies to demonstrate that dietary context significantly alters the hepatic transcriptomic and metabolic responses to a pesticide mixture in a diet specific manner. The differential response observed between CD- and WD-fed mice emphasizes the complex interplay between environmental contaminants and dietary factors and the importance of considering dietary context when evaluating the metabolic effects of pesticide mixtures. Despite the limitations in translating findings from mice to humans, our results suggest that sensitivity to pesticide exposure may differ according to metabolic status. Given that toxicological reference values ensuring consumer safety are defined for individual pesticides, our findings suggest that their relevance may differ when pesticides are combined in mixtures.

### Limitations of the study

Although our study assessed the effects of a pesticide mixture administered through food intake, at two toxicological reference doses, on 12 animals per group and in two nutritional contexts, it has some limitations. First, we did not identify the mechanisms underlying the effects of the pesticide mixture on glucose homeostasis or the specific tissue in which pesticides accumulate in WD-fed mice. However, our study provides a comprehensive overview of the observed transcriptomic and phenotypic outcomes, which can serve as a basis for future mechanistic studies. Further mouse studies comparing the effects of the pesticide mixture with those of individual pesticides would be necessary to provide a better understanding of the interactions among the compounds in the mixture. Finally, we did not evaluate the effects of pesticide exposure in female mice. As energy and xenobiotic metabolism in the liver are highly sexually dimorphic, it is likely that the pesticide cocktail could have sex-specific health effects.

## Acknowledgments

This work was supported by the French Foundation for the Medical Research FRM (ENV202109013962), the Caisse Centrale de la Mutualité Sociale Agricole (AAP Mutualité Sociale Agricole MSA 2022-BIOMEC), the department AlimH of INRAE. We thank Professor Bart Staels and Dr N. Hennuyer (Institut Pasteur, Lille, France) for their generous gift of IHH cells. We thank the EZOP staff, the GeT-Trix Genotoul facility, Metatoul-Metabohub, Anexplo, and We-Met facilities for their help. We also thank the INRAE SAAJ–RAF team (Jouy-en-Josas, France) for its technical support with the pellet preparation. C.R. was supported by FRM (ENV202109013962).

## References

Ahmad MF, Ahmad FA, Alsayegh AA, Zeyaullah Md, AlShahrani AM, Muzammil K, et al. 2024. Pesticides impacts on human health and the environment with their mechanisms of action and possible countermeasures. Heliyon 10:e29128; doi:10.1016/j.heliyon.2024.e29128.

Akpolat N, Yahsi S, Godekmerdan A, Yalniz M, Demirbag K. 2005. The value of alpha-SMA in the evaluation of hepatic fibrosis severity in hepatitis B infection and cirrhosis development: a histopathological and immunohistochemical study. Histopathology 47:276–280; doi:10.1111/j.1365-2559.2005.02226.x.

Arab A, Mostafalou S. 2023. Pesticides and insulin resistance-related metabolic diseases: Evidences and mechanisms. Pesticide Biochemistry and Physiology 195:105521; doi:10.1016/j.pestbp.2023.105521.

Arciello M, Gori M, Maggio R, Barbaro B, Tarocchi M, Galli A, et al. 2013. Environmental Pollution: A Tangible Risk for NAFLD Pathogenesis. IJMS 14:22052–22066; doi:10.3390/ijms141122052.

Attema B, Janssen AWF, Rijkers D, van Schothorst EM, Hooiveld GJEJ, Kersten S. 2022. Exposure to low-dose perfluorooctanoic acid promotes hepatic steatosis and disrupts the hepatic transcriptome in mice. Molecular Metabolism 66:101602; doi:10.1016/j.molmet.2022.101602.

Barrios-Rodríguez R, Pérez-Carrascosa FM, Gómez-Peña C, Mustieles V, Salcedo-Bellido I, Requena P, et al. 2021. Associations of accumulated selected persistent organic pollutants in adipose tissue with insulin sensitivity and risk of incident type-2 diabetes. Environment International 155:106607; doi:10.1016/j.envint.2021.106607.

Baudry J, Lelong H, Adriouch S, Julia C, Allès B, Hercberg S, et al. 2018. Association between organic food consumption and metabolic syndrome: cross-sectional results from the NutriNet-Santé study. Eur J Nutr 57:2477–2488; doi:10.1007/s00394-017-1520-1.

Baudry J, Rebouillat P, Allès B, Cravedi J-P, Touvier M, Hercberg S, et al. 2021. Estimated dietary exposure to pesticide residues based on organic and conventional data in omnivores, pesco-vegetarians, vegetarians and vegans. Food and Chemical Toxicology 153:112179; doi:10.1016/j.fct.2021.112179.

Capitão A, Lyssimachou A, Castro LFC, Santos MM. 2017. Obesogens in the aquatic environment: an evolutionary and toxicological perspective. Environment International 106:153–169; doi:10.1016/j.envint.2017.06.003.

Castorina R, Bradman A, McKone TE, Barr DB, Harnly ME, Eskenazi B. 2003. Cumulative organophosphate pesticide exposure and risk assessment among pregnant women living in an agricultural community: a case study from the CHAMACOS cohort. Environ Health Perspect 111:1640–1648; doi:10.1289/ehp.5887.

Cedergreen N. 2014. Quantifying Synergy: A Systematic Review of Mixture Toxicity Studies within Environmental Toxicology. A. Nazir, ed PLoS ONE 9:e96580; doi:10.1371/journal.pone.0096580.

Chang J, Li J, Wang H, Wang Y, Guo B, Yin J, et al. 2016. Tissue distribution, metabolism and hepatic tissue injury in Chinese lizards (Eremias argus) after a single oral administration of lambda-cyhalothrin. Environmental Pollution 218:965–972; doi:10.1016/j.envpol.2016.08.045.

Chousidis I, Ellero-Simatos S, Payrastre L, Payros G, Lippi Y, Blaquière M, et al. 2025. Bisphenol analogue exposure at low concentrations modifies heart-brain functions and transcriptomics in zebrafish larvae. Journal of Hazardous Materials 498:140004; doi:10.1016/j.jhazmat.2025.140004.

Christen V, Crettaz P, Fent K. 2014. Additive and synergistic antiandrogenic activities of mixtures of azol fungicides and vinclozolin. Toxicology and Applied Pharmacology 279:455–466; doi:10.1016/j.taap.2014.06.025.

Costes S, Bertrand G, Ravier MA. 2021. Mechanisms of Beta-Cell Apoptosis in Type 2 Diabetes-Prone Situations and Potential Protection by GLP-1-Based Therapies. IJMS 22:5303; doi:10.3390/ijms22105303.

de Sousa G, Nawaz A, Cravedi J-P, Rahmani R. 2014. A Concentration Addition Model to Assess Activation of the Pregnane X Receptor (PXR) by Pesticide Mixtures Found in the French Diet. Toxicological Sciences 141:234–243; doi:10.1093/toxsci/kfu120.

Di Tommaso P, Chatzou M, Floden EW, Barja PP, Palumbo E, Notredame C. 2017. Nextflow enables reproducible computational workflows. Nat Biotechnol 35:316–319; doi:10.1038/nbt.3820.

Dinca V, Docea AO, Drocas AI, Nikolouzakis TK, Stivaktakis PD, Nikitovic D, et al. 2023. A mixture of 13 pesticides, contaminants, and food additives below individual NOAELs produces histopathological and organ weight changes in rats. Arch Toxicol 97:1285–1298; doi:10.1007/s00204-023-03455-x.

Djekkoun N, Lalau J-D, Bach V, Depeint F, Khorsi-Cauet H. 2021. Chronic oral exposure to pesticides and their consequences on metabolic regulation: role of the microbiota. Eur J Nutr 60:4131–4149; doi:10.1007/s00394-021-02548-6.

Docea AO, Gofita E, Goumenou M, Calina D, Rogoveanu O, Varut M, et al. 2018. Six months exposure to a real life mixture of 13 chemicals’ below individual NOAELs induced non monotonic sex-dependent biochemical and redox status changes in rats. Food and Chemical Toxicology 115:470–481; doi:10.1016/j.fct.2018.03.052.

Docea AO, Goumenou M, Calina D, Arsene AL, Dragoi CM, Gofita E, et al. 2019. Adverse and hormetic effects in rats exposed for 12 months to low dose mixture of 13 chemicals: RLRS part III. Toxicology Letters 310:70–91; doi:10.1016/j.toxlet.2019.04.005.

European Food Safety Authority (EFSA), Carrasco Cabrera L, Di Piazza G, Dujardin B, Marchese E, Medina Pastor P. 2024. The 2022 European Union report on pesticide residues in food. EFS2 22; doi:10.2903/j.efsa.2024.8753.

Expertise Collective Inserm. 2013. Pesticides: Effets sur la santé. Les éditions Inserm. Paris.

Expertise Collective Inserm. 2021. Pesticides et effets sur la santé. Nouvelles données. Editions EDP science.

Fountoucidou P, Veskoukis AS, Kerasioti E, Docea AO, Taitzoglou IA, Liesivuori J, et al. 2019. A mixture of routinely encountered xenobiotics induces both redox adaptations and perturbations in blood and tissues of rats after a long-term low-dose exposure regimen: The time and dose issue. Toxicology Letters 317:24–44; doi:10.1016/j.toxlet.2019.09.015.

Friedman SL, Neuschwander-Tetri BA, Rinella M, Sanyal AJ. 2018. Mechanisms of NAFLD development and therapeutic strategies. Nat Med 24:908–922; doi:10.1038/s41591-018-0104-9.

Fujino C, Watanabe Y, Sanoh S, Nakajima H, Uramaru N, Kojima H, et al. 2019. Activation of PXR, CAR and PPARα by pyrethroid pesticides and the effect of metabolism by rat liver microsomes. Heliyon 5:e02466; doi:10.1016/j.heliyon.2019.e02466.

Groswald AM, Gripshover TC, Watson WH, Wahlang B, Luo J, Jophlin LL, et al. 2023. Investigating the Acute Metabolic Effects of the N-Methyl Carbamate Insecticide, Methomyl, on Mouse Liver. Metabolites 13:901; doi:10.3390/metabo13080901.

He B, Ni Y, Jin Y, Fu Z. 2020. Pesticides-induced energy metabolic disorders. Science of The Total Environment 729:139033; doi:10.1016/j.scitotenv.2020.139033.

Hectors TLM, Vanparys C, van der Ven K, Martens GA, Jorens PG, Van Gaal LF, et al. 2011. Environmental pollutants and type 2 diabetes: a review of mechanisms that can disrupt beta cell function. Diabetologia 54:1273–1290; doi:10.1007/s00125-011-2109-5.

Hernández AF, Parrón T, Tsatsakis AM, Requena M, Alarcón R, López-Guarnido O. 2013. Toxic effects of pesticide mixtures at a molecular level: Their relevance to human health. Toxicology 307:136–145; doi:10.1016/j.tox.2012.06.009.

Hoyeck MP, Matteo G, MacFarlane EM, Perera I, Bruin JE. 2022. Persistent organic pollutants and β-cell toxicity: a comprehensive review. American Journal of Physiology-Endocrinology and Metabolism 322:E383–E413; doi:10.1152/ajpendo.00358.2021.

Jabłońska-Trypuć. 2017. The impact of pesticides on oxidative stress level in human organism and their activity as an endocrine disruptor. Journal of Environmental Science and Health, Part B.

Jackson E, Shoemaker R, Larian N, Cassis L. 2017. Adipose Tissue as a Site of Toxin Accumulation. In: Comprehensive Physiology (R. Terjung, ed). Wiley. 1085–1135.

Kesse-Guyot E, Baudry J, Assmann KE, Galan P, Hercberg S, Lairon D. 2017. Prospective association between consumption frequency of organic food and body weight change, risk of overweight or obesity: results from the NutriNet-Santé Study. Br J Nutr 117:325–334; doi:10.1017/s0007114517000058.

Kesse-Guyot E, Rebouillat P, Payrastre L, Allès B, Fezeu LK, Druesne-Pecollo N, et al. 2020. Prospective association between organic food consumption and the risk of type 2 diabetes: findings from the NutriNet-Santé cohort study. Int J Behav Nutr Phys Act 17; doi:10.1186/s12966-020-01038-y.

Knebel C, Kebben J, Eberini I, Palazzolo L, Hammer HS, Süssmuth RD, et al. 2018a. Propiconazole is an activator of AHR and causes concentration additive effects with an established AHR ligand. Arch Toxicol 92:3471–3486; doi:10.1007/s00204-018-2321-x.

Knebel C, Kebben J, Eberini I, Palazzolo L, Hammer HS, Süssmuth RD, et al. 2018b. Propiconazole is an activator of AHR and causes concentration additive effects with an established AHR ligand. Arch Toxicol 92:3471–3486; doi:10.1007/s00204-018-2321-x.

Lee H, Gao Y, Kim JK, Shin S, Choi M, Hwang Y, et al. 2023. Synergetic effects of concurrent chronic exposure to a mixture of OCPs and high-fat diets on type 2 diabetes and beneficial effects of caloric restriction in female zebrafish. Journal of Hazardous Materials 446:130659; doi:10.1016/j.jhazmat.2022.130659.

Lee Y-M, Ha C-M, Kim S-A, Thoudam T, Yoon Y-R, Kim D-J, et al. 2017. Low-Dose Persistent Organic Pollutants Impair Insulin Secretory Function of Pancreatic b-Cells: Human and In Vitro Evidence. 66.

Léger T, Balaguer P, Le Hégarat L, Fessard V. 2023. Fate and PPARγ and STATs-driven effects of the mitochondrial complex I inhibitor tebufenpyrad in liver cells revealed with multi-omics. Journal of Hazardous Materials 442:130083; doi:10.1016/j.jhazmat.2022.130083.

Leroux P. 2003. Modes d’action des produits phytosanitaires sur les organismes pathogènes des plantes. Comptes Rendus Biologies 326:9–21; doi:10.1016/S1631-0691(03)00005-2.

Li J, Li X, Zhang Z, Cheng W, Liu G, Zhao G. 2023. High-Fat Diet Aggravates the Disorder of Glucose Metabolism Caused by Chlorpyrifos Exposure in Experimental Rats. Foods 12:816; doi:10.3390/foods12040816.

Li W, Xiao H, Wu H, Pan C, Deng K, Xu X, et al. 2022. Analysis of environmental chemical mixtures and nonalcoholic fatty liver disease: NHANES 1999–2014. Environmental Pollution 311:119915; doi:10.1016/j.envpol.2022.119915.

Lichtenstein D, Luckert C, Alarcan J, de Sousa G, Gioutlakis M, Katsanou ES, et al. 2020. An adverse outcome pathway-based approach to assess steatotic mixture effects of hepatotoxic pesticides in vitro. Food and Chemical Toxicology 139:111283; doi:10.1016/j.fct.2020.111283.

Lippi Y, Soubès F. 2023. MATRiX: a shiny application for Mining and functional Analysis of TRanscriptomics data.; doi:10.17180/DJ0G-5F33.

Lukowicz C, Ellero-Simatos S, Régnier M, Polizzi A, Lasserre F, Montagner A, et al. 2018. Metabolic Effects of a Chronic Dietary Exposure to a Low-Dose Pesticide Cocktail in Mice: Sexual Dimorphism and Role of the Constitutive Androstane Receptor. Environ Health Perspect 126:067007; doi:10.1289/EHP2877.

Ma Q, Deng P, Lin M, Yang L, Li L, Guo L, et al. 2021. Long-term bisphenol A exposure exacerbates diet-induced prediabetes via TLR4-dependent hypothalamic inflammation. Journal of Hazardous Materials 402:123926; doi:10.1016/j.jhazmat.2020.123926.

Mardinoglu A. 2018. Systems biology in hepatology: approaches and applications.

Mellor CL, Steinmetz FP, Cronin MTD. 2016. The identification of nuclear receptors associated with hepatic steatosis to develop and extend adverse outcome pathways. Critical Reviews in Toxicology 46:138–152; doi:10.3109/10408444.2015.1089471.

Mesnage R, Mahmud N, Mein CA, Antoniou MN. 2021. Alterations in small RNA profiles in liver following a subchronic exposure to a low-dose pesticide mixture in Sprague-Dawley rats. Toxicology Letters 353:20–26; doi:10.1016/j.toxlet.2021.10.001.

Nichols RG, Rimal B, Hao F, Peters JM, Davenport ER, Patterson AD. 2024. Chlorpyrifos modulates the mouse gut microbiota and metabolic activity. Environment International 192:109022; doi:10.1016/j.envint.2024.109022.

Podechard N, Ducheix S, Polizzi A, Lasserre F, Montagner A, Legagneux V, et al. 2018. Dual extraction of mRNA and lipids from a single biological sample. Sci Rep 8:7019; doi:10.1038/s41598-018-25332-9.

Rajak, S., Raza, z, Tewari, A. 2022. Environmental Toxicants and NAFLD: A Neglected yet Significant Relationship. Dig Dis Sci.

Rebouillat P, Vidal R, Cravedi J-P, Taupier-Letage B, Debrauwer L, Gamet-Payrastre L, et al. 2021. Estimated dietary pesticide exposure from plant-based foods using NMF-derived profiles in a large sample of French adults. Eur J Nutr 60:1475–1488; doi:10.1007/s00394-020-02344-8.

Rebouillat P, Vidal R, Cravedi J-P, Taupier-Letage B, Debrauwer L, Gamet-Payrastre L, et al. 2022. Prospective association between dietary pesticide exposure profiles and type 2 diabetes risk in the NutriNet-Santé cohort. Environ Health 21:57; doi:10.1186/s12940-022-00862-y.

Rives C, Fougerat A, Ellero-Simatos S, Loiseau N, Guillou H, Gamet-Payrastre L, et al. 2020. Oxidative Stress in NAFLD: Role of Nutrients and Food Contaminants. Biomolecules 10:1702; doi:10.3390/biom10121702.

Roustan A, Aye M, De Meo M, Di Giorgio C. 2014. Genotoxicity of mixtures of glyphosate and atrazine and their environmental transformation products before and after photoactivation. Chemosphere 108:93–100; doi:10.1016/j.chemosphere.2014.02.079.

Schmidt FF, Lichtenstein D, Planatscher H, Mentz A, Kalinowski J, Steinhilber AE, et al. 2021. Pesticide mixture effects on liver protein abundance in HepaRG cells. Toxicology 458:152839; doi:10.1016/j.tox.2021.152839.

Sousa S, Rede D, Cruz Fernandes V, Pestana D, Faria G, Delerue-Matos C, et al. 2023. Accumulation of organophosphorus pollutants in adipose tissue of obese women - metabolic alterations. Environmental Research 239:117337; doi:10.1016/j.envres.2023.117337.

Tait S, Lori G, Tassinari R, La Rocca C, Maranghi F. 2022. In Vitro Assessment and Toxicological Prioritization of Pesticide Mixtures at Concentrations Derived from Real Exposure in Occupational Scenarios. IJERPH 19:5202; doi:10.3390/ijerph19095202.

Velmurugan G, Ramprasath T, Swaminathan K, Mithieux G, Rajendhran J, Dhivakar M, et al. 2017. Gut microbial degradation of organophosphate insecticides-induces glucose intolerance via gluconeogenesis. Genome Biol 18:8; doi:10.1186/s13059-016-1134-6.

Vinken M, Knapen D, Vergauwen L, Hengstler JG, Angrish M, Whelan M. 2017. Adverse outcome pathways: a concise introduction for toxicologists. Arch Toxicol 91:3697–3707; doi:10.1007/s00204-017-2020-z.

Wahlang B, Jin J, Beier JI, Hardesty JE, Daly EF, Schnegelberger RD, et al. 2019. Mechanisms of Environmental Contributions to Fatty Liver Disease. Curr Envir Health Rpt 6:80–94; doi:10.1007/s40572-019-00232-w.

Wang B, Tsakiridis EE, Zhang S, Llanos A, Desjardins EM, Yabut JM, et al. 2021. The pesticide chlorpyrifos promotes obesity by inhibiting diet-induced thermogenesis in brown adipose tissue. Nat Commun 12:5163; doi:10.1038/s41467-021-25384-y.

Wang R, Yang X, Wang T, Kou R, Liu P, Huang Y, et al. 2023. Synergistic effects on oxidative stress, apoptosis and necrosis resulting from combined toxicity of three commonly used pesticides on HepG2 cells. Ecotoxicology and Environmental Safety 263:115237; doi:10.1016/j.ecoenv.2023.115237.

Wang X, Lu Q, Guo J, Ares I, Martínez M, Martínez-Larrañaga M-R, et al. 2022. Oxidative Stress and Metabolism: A Mechanistic Insight for Glyphosate Toxicology. Annu Rev Pharmacol Toxicol 62:617–639; doi:10.1146/annurev-pharmtox-020821-111552.

Wei Y, Liu W, Liu J. 2023a. Environmentally relevant exposure to cypermethrin aggravates diet-induced diabetic symptoms in mice: The interaction between environmental chemicals and diet. Environment International 178:108090; doi:10.1016/j.envint.2023.108090.

Wei Y, Wang L, Liu J. 2023b. The diabetogenic effects of pesticides: Evidence based on epidemiological and toxicological studies. Environmental Pollution 331:121927; doi:10.1016/j.envpol.2023.121927.

Xiao X, Sun Q, Kim Y, Yang S-H, Qi W, Kim D, et al. 2018. Exposure to permethrin promotes high fat diet-induced weight gain and insulin resistance in male C57BL/6J mice. Food and Chemical Toxicology 111:405–416; doi:10.1016/j.fct.2017.11.047.

Yang D, Sun X, Wei X, Zhang B, Fan X, Du H, et al. 2023. Lambda-cyhalothrin induces lipid accumulation in mouse liver is associated with AMPK inactivation. Food and Chemical Toxicology 172:113563; doi:10.1016/j.fct.2022.113563.

Yang JS, Park Y. 2018. Insecticide Exposure and Development of Nonalcoholic Fatty Liver Disease. J Agric Food Chem 66:10132–10138; doi:10.1021/acs.jafc.8b03177.

Zhou Y, Zhou B, Pache L, Chang M, Khodabakhshi AH, Tanaseichuk O, et al. 2019. Metascape provides a biologist-oriented resource for the analysis of systems-level datasets. Nat Commun 10:1523; doi:10.1038/s41467-019-09234-6.

