## supplementary material for "Diet modulates metabolic and hepatic responses to chronic pesticide mixture exposure in mice"

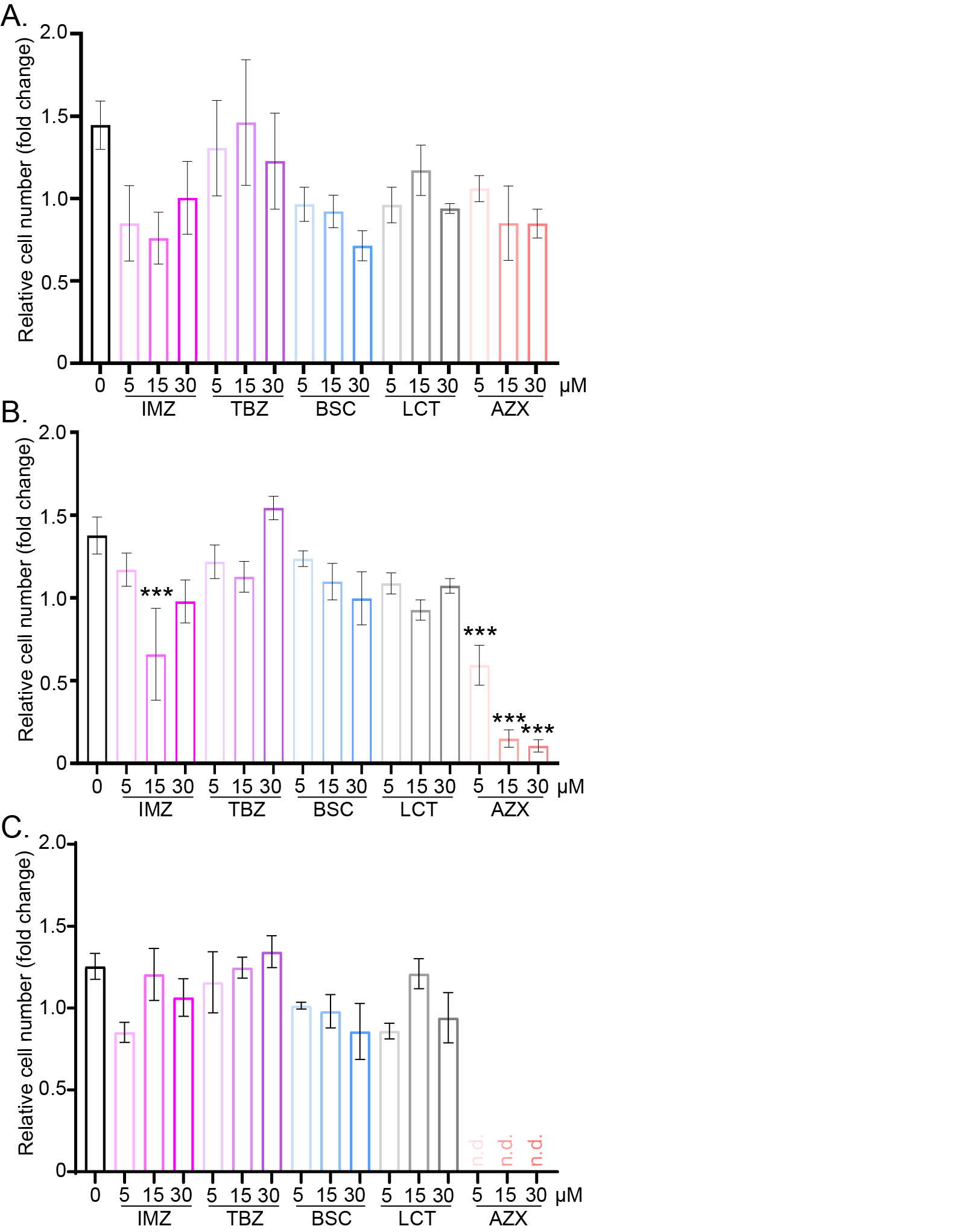

**Supplementary Figure 1:** Cell viability of IHH cells treated with individual pesticides IMZ, TBZ, BSC, and LCT at 5, 15, 30 µM for 24 h (**A**), 72 h (**B**) or 10 days (**C**) (n = 3). Data were normalized to the total mean of each experiment before being pooled. Data are presented as the mean ± SEM (n.d., not-detected). *Treatment effect, **p* < 0.05, ***p* < 0.01, ****p* < 0.001 (one-way ANOVA followed by Tuckey’s post-hoc test).

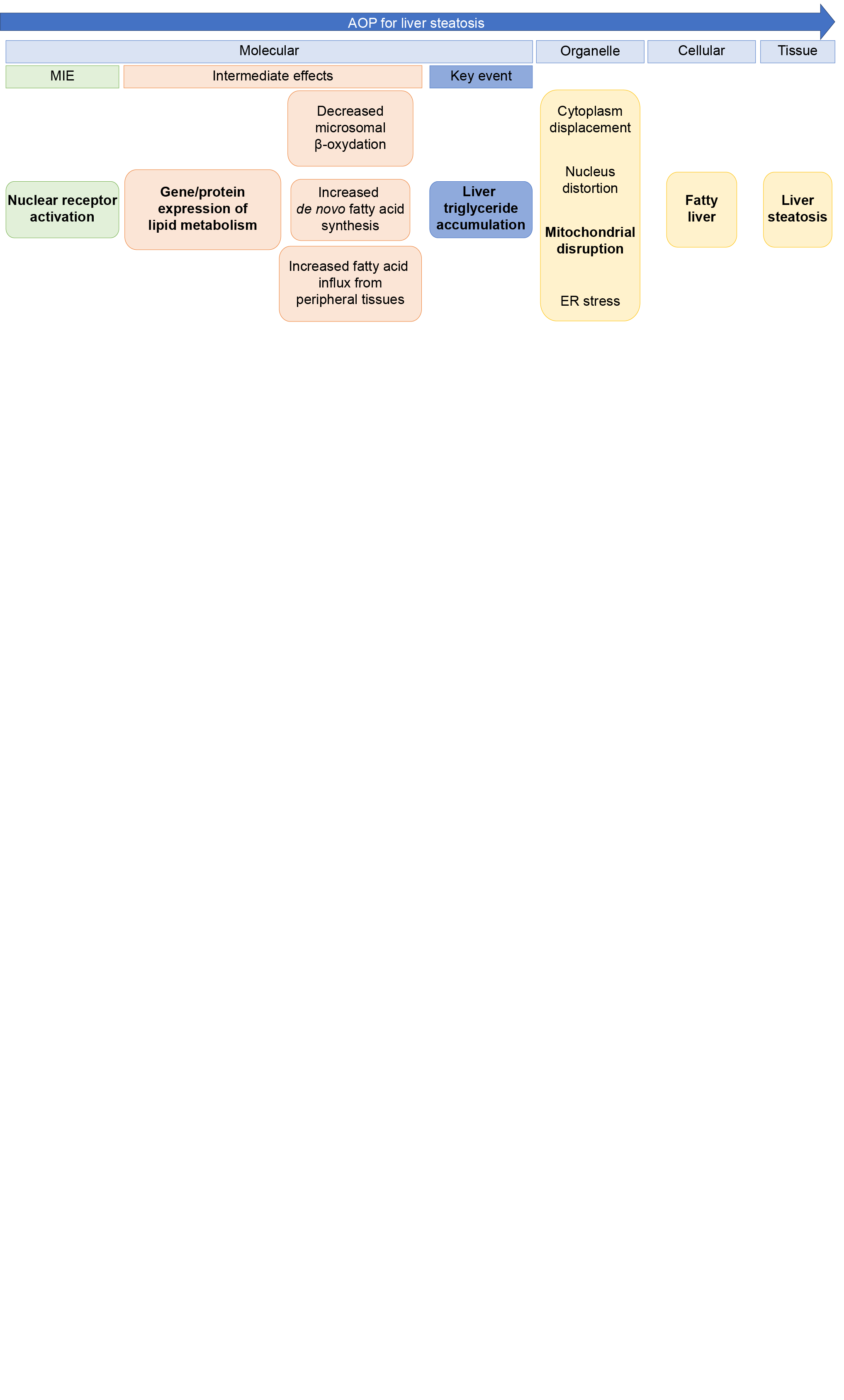

**Supplementary Figure 2:** Adverse outcome pathway (AOP) for liver steatosis (adapted from Mellor et al. 2016 and Vinken et al. 2017). *In vitro* end points analyzed in this study are shown in bold. MIE: molecular initiating events.

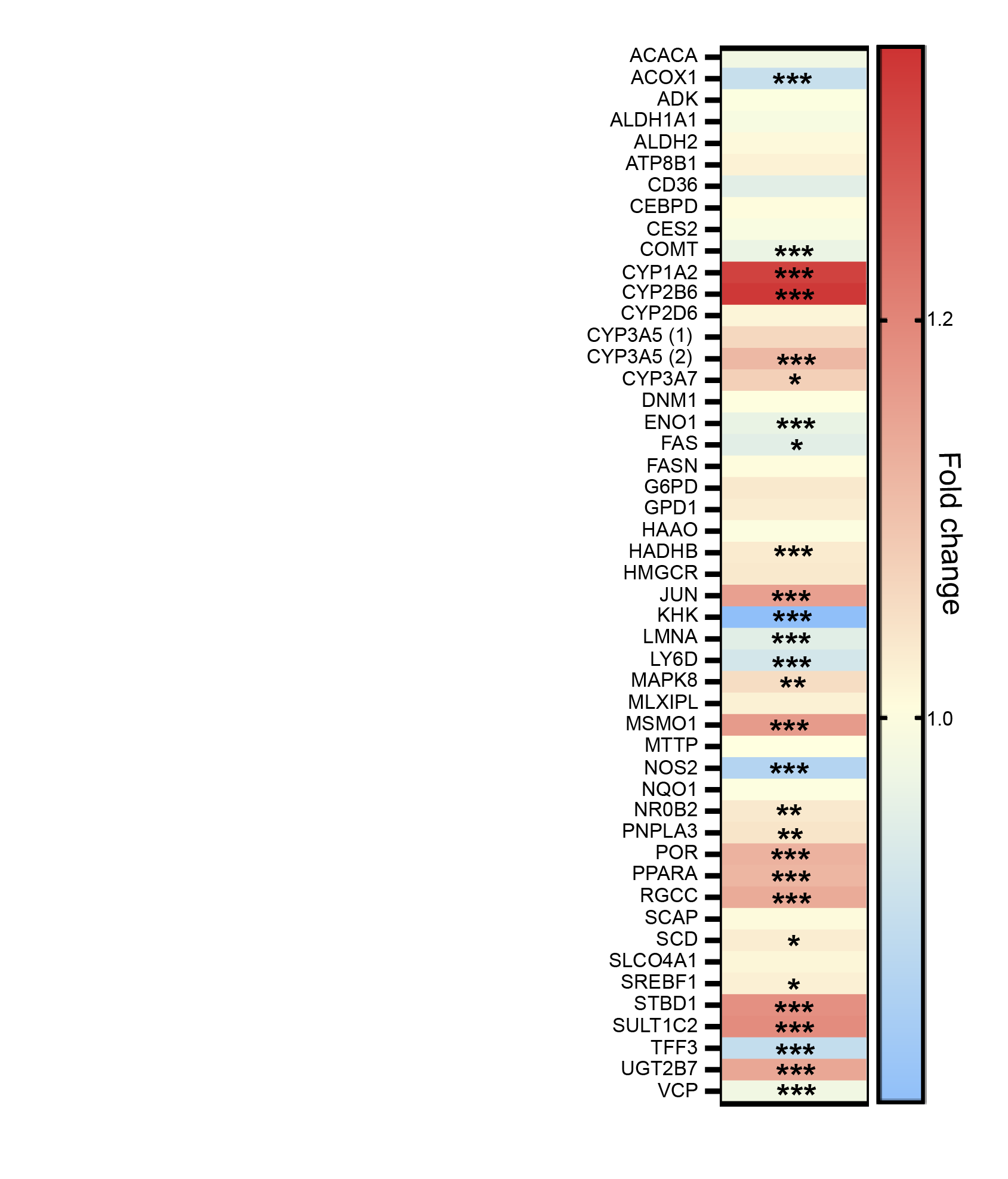

**Supplementary Figure 3:** Heatmap representing the log(fold-change) of gene expression between IHH cells treated with pesticide mixture (ITBL) and untreated cells for a set of 48 genes linked to liver steatosis, hepatotoxicity, and nuclear receptor activation (Lichtenstein et al. 2020). *Treatment effect, **p* < 0.05, ***p* < 0.01, ****p* < 0.001 (one-way ANOVA followed by Tukey's post-hoc test).

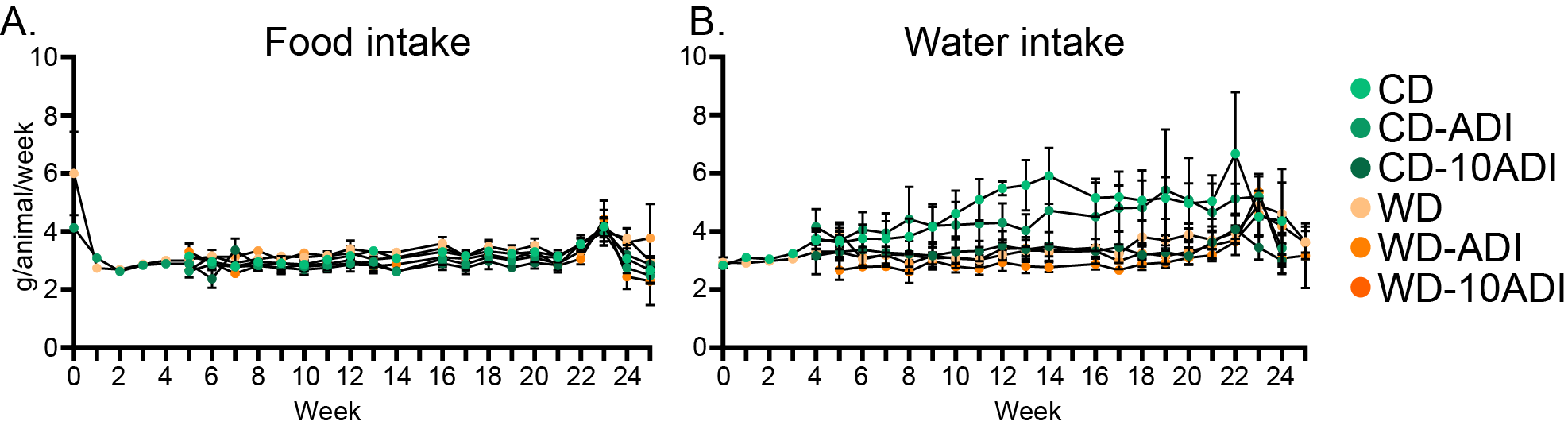

**Supplementary Figure 4:** Food (**A**) and water (**B**) intake per animal and per week in each group of mice from week 1 through 25 weeks (n = 12 per group). Data are presented as the mean ± SEM.

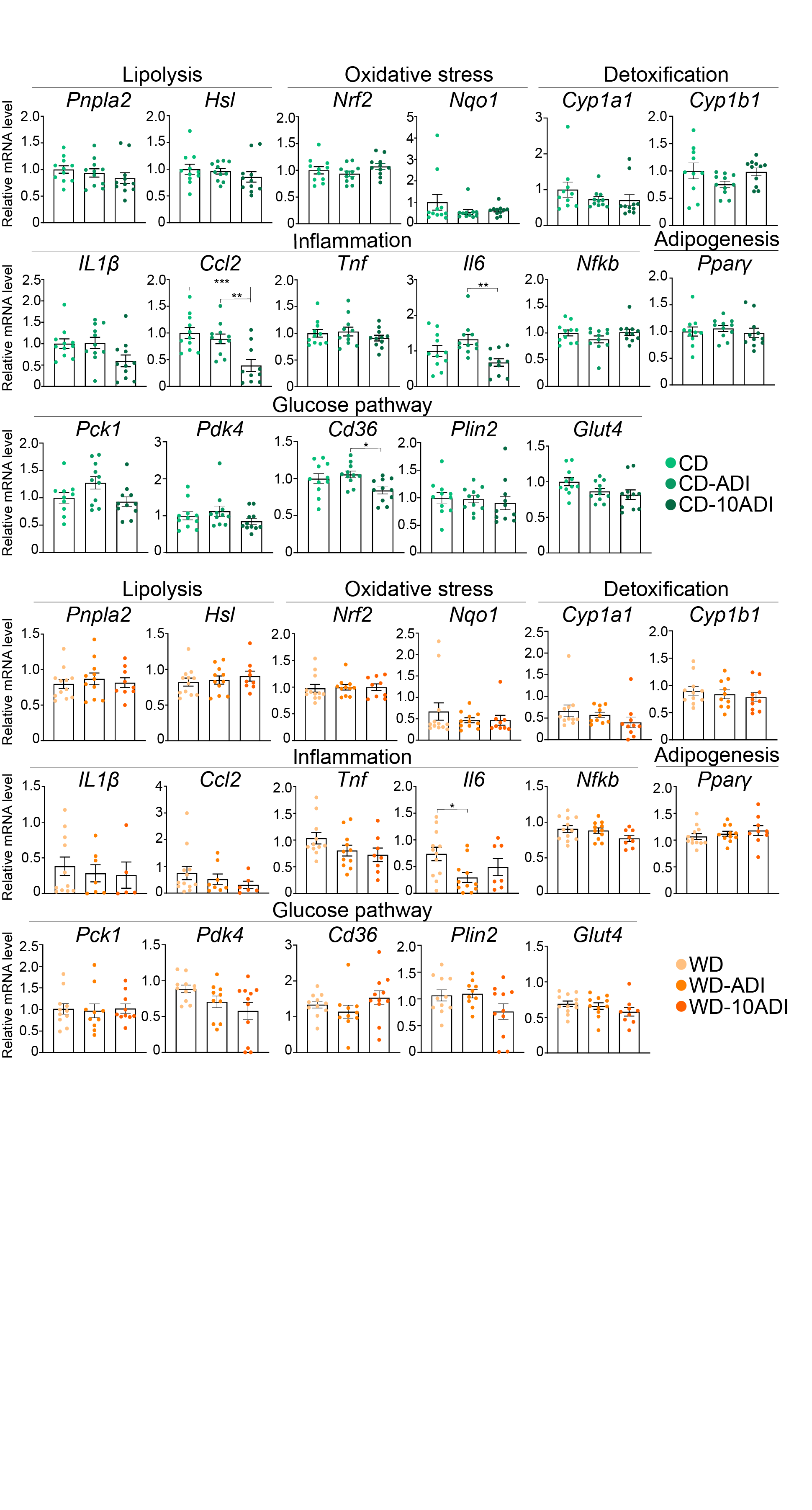

**Supplementary Figure 5:** mRNA expression of genes involved in lipolysis, oxidative stress, adipogenesis, glucose metabolism, inflammation and in xenobiotic metabolism measured by RT-qPCR in white adipose tissue samples from each group of CD- and WD-fed mice (n = 12 per group). Data are presented as the mean ± SEM. * exposed vs. non-exposed mice, **p* < 0.05, ***p* < 0.01, ****p* < 0.001 (two-way ANOVA followed by Tukey's post-hoc test).

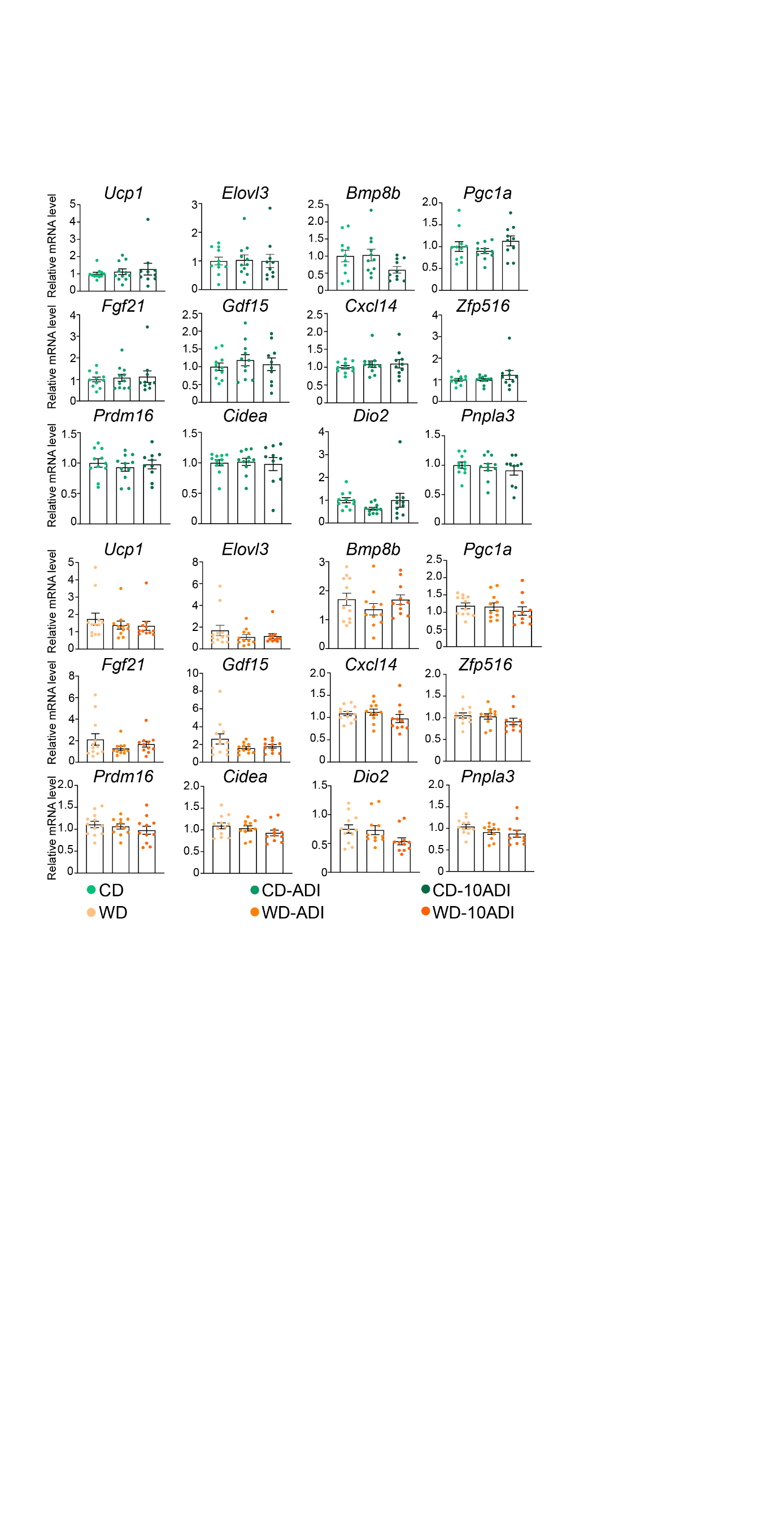

**Supplementary Figure 6:** mRNA expression of genes involved in brown adipose tissue activation and encoding batokines measured by RT-qPCR in brown adipose tissue samples from each group of CD- and WD-fed mice (n = 12 per group). Data are presented as the mean ± SEM. *Exposed vs. non-exposed mice, **p* < 0.05, ***p* < 0.01, ****p* < 0.001 (two-way ANOVA followed by Tukey's post-hoc test).

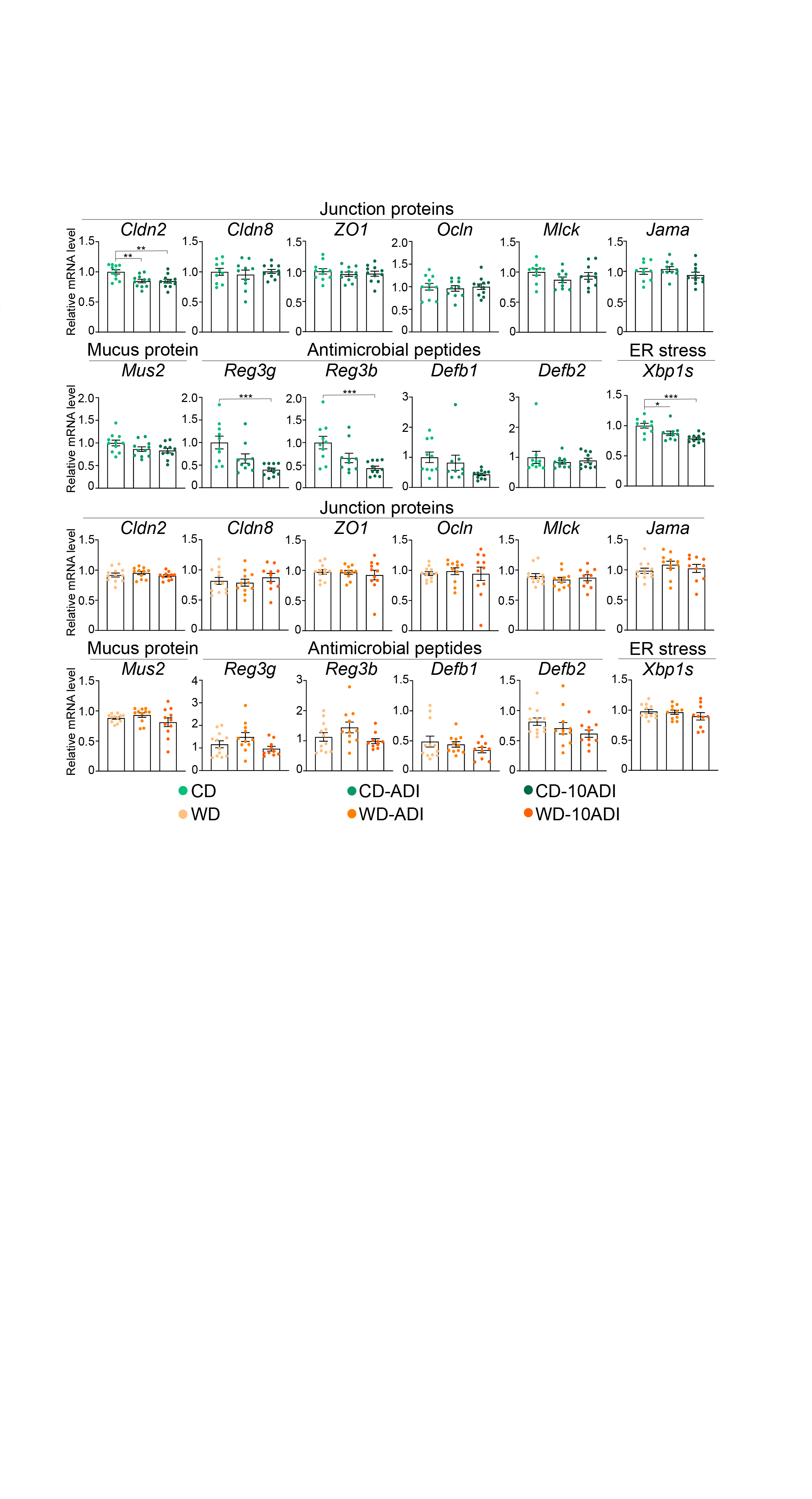

**Supplementary Figure 7:** mRNA expression of genes encoding junction proteins, mucus proteins, antimicrobial peptides and proteins involved in ER stress measured by RT-qPCR in ileum samples from each group of CD- and WD-fed mice (n = 12 per group). Data are presented as the mean ± SEM. *Exposed vs. non-exposed mice, **p* < 0.05, ***p* < 0.01, ****p* < 0.001 (two-way ANOVA followed by Tukey's post-hoc test).

**Supplementary Table 1:** Chemical families, functions, and acceptable daily intake (ADI, mg/kg body weight/day) of each pesticide and the expected and measured pesticide concentrations (determined level, µg/kg food) in the animal pellets.

| **Pesticide** | **Chemical family** | **Function** | **ADI**  **(mg/kg BW/day)** | **Expected quantity (µg/kg food)**  **ADI** | **Expected quantity (µg/kg food)**  **10ADI** | **Determined level**  **(µg/kg food)** | | | | |
| --- | --- | --- | --- | --- | --- | --- | --- | --- | --- | --- |
|  |  |  |  |  |  | **CD ADI** | **WD ADI** | **CD 10ADI** | **WD 10ADI** | |
| **Boscalid** | **Carboxamide** | **Fungicide** | **0.04** | **200** | **2000** | **200 ± 100** | **≤ 300** | **2800 ± 1400** | | **1500 ± 80** |
| **Imazalil** | **Imidazole** | **Fungicide** | **0.025** | **125** | **1250** | **48 ± 24** | **120 ± 60** | **990 ± 500** | | **1200 ± 600** |
| **Thiabendazole** | **Benzimidazole** | **Fungicide** | **0.1** | **500** | **5000** | **640 ± 320** | **780 ± 390** | **5200 ±2600** | | **5000 ± 2500** |
| **Lambda−cyhalothrin** | **Pyrethroid** | **Insecticide** | **0.0025** | **12.5** | **125** | **≤ LOQ** | **≤ LOQ** | **≤ LOQ** | | **120 ± 60** |

Note: CD, chow diet; WD, western diet; BW, body weight; ADI, acceptable daily intake (<https://ephy.anses.fr/>). Expected quantity refers to the incorporated quantities of pesticides in food pellets.

**Supplementary Table 2:** Sequences of primers used in qPCR.

| Genes | Forward primers | Reverse primers |
| --- | --- | --- |
| GAPDH | GGGAAGGTGAAGGTCGGAGT | TGACCAGGCGCCCAATAC |
| CYP4A11 | GGGCTCATGGCAGACTCTG | GGCCAAGGAGCTCTTCCC |
| CYP1A1 | CTACAGCTTCATGCAGAAGATGGT | TCTGTGATGTCCCGGATGTG |
| CYP2B6 | GGAGGAGCGGATTCAGGAG | TGGAATCGTTTTCCAAAGACG |
| CYP3A4 | GATGGTCAACAGCCTGTGCT | CCTCCGGTTTGTGAAGACAGAA |
| SREBP1c | GGCGAGCCGTGCGAT | TGCCCCACTCCCAGCAT |
| Pnpla2 | AGTGTCCTTCACCATCCGCTT | GGATATCTTCAGGGACATCAGGC |
| Hsl | GGCTTACTGGGCACAGATACCT | CTGAAGGCTCTGAGTTGCTCAA |
| Pparγ | GATGCACTGCCTATGAGCACTT | GAATGGCATCTCTGTGTCAACC |
| Glut4 | CCAGCAGATCGGCTCTGAC | GCCAAGCACAGCTGAGAATACA |
| Nrf2 | TTGCCACCGCCAGGACTA | TGTCTTGCCTCCAAAGGATGT |
| Nqo1 | GGGACATGAACGTCATTCTCTG | GTTCTAAGACCTGGAAGCCACAG |
| Il1β | GCCCATCCTCTGTGACTCAT | AGGCCACAGGTATTTTGTCG |
| Ccl2 | GGTGTCCCAAAGAAGCTGTAGTTTT | AGTTGTAGGTTCTGATCTCATTTGGTT |
| Tnf | TCCCCAAAGGGATGAGAAGTTC | GCGCTGGCTCAGCCACT |
| Il6 | AGCCAGAGTCCTTCAGAGAGATACA | TTGGTCCTTAGCCACTCCTTCT |
| Nfkb | GACGATGATCCCTACGGAACTG | CTGCATTAAATATTGAGTGAGTCAAAGC |
| Cyp1a1 | CACTACAGGACATTTGAGAAGGGC | GCTCAATGAGGCTGTCTGTGAT |
| Cyp1b1 | TGTGGCTGCTCATCCTCTTTAC | CCCCACAACCTGGTCCAAC |
| Pck1 | GAACCCCAGCCTGCCC | GAGCAACTCCAAAAAACCCG |
| Pdk4 | ATCGCCAGAATTAAACCTCACAC | TGGATTGGTTGGCCTGGA |
| Cd36 | GTGTTTGGAGGCATTCT | GAAAGCAGTGGTTCCTT |
| Plin2 | CCATTTCTCAGCTCCACTCCAC | GTGTCGTCGTAGCCGATGC |
| Ucp1 | CCTGCCTCTCTCGGAAACAA | TGTAGGCTGCCCAATGAACA |
| Elovl3 | GCCTCTCATCCTCTGGTCCT | TGCCATAAACTTCCACATCCT |
| Bmp8b | GTGCTGACCTGATTATGAGCTTT | GGTACGGTCGCGTTCCACTA |
| Pgc1α | CAATCGGAAATCATATCCAACCA | CTGTGAGGACCGCTAGGAAGT |
| Fgf21 | AAAGCCTCTAGGTTTCTTTGCCA | CCTCAGGATCAAAGTGAGGCG |
| Gfd15 | AGCCGAGAGGACTCGAACTCA | TGGGACCCCAATCTCACCTCT |
| Cxcl14 | TACCCACACTGCGAGGAGAA | GGACATGCTCTTGGTGGTGA |
| Prdm16 | AAGGCGAGGGCGAGGAA | CATATTATTTACAACGTCACCGTCACT |
| Cidea | CTACGCGGGAGCCCTCA | GGGCGAGCTGGATGTATGA |
| Dio2 | AGCTTCCTCCTAGATGCCTACA | GGGAGCATCTTCACCCAGTTTA |
| Pnpla3 | ACGCGGTCACCTTCGTGT | AGCCCGTCTCTGATGCACTT |
| Zfp516 | AGTGGTGAAGAGGCTGTGCCTGAA | AAGAGCAGCAGTGGCGAGGCT |
| Cldn2 | TCTCAGCCCTGTTTTCTTTGGT | GGGCCTGGTAGCCATCATAGTA |
| Cldn8 | GGAGGAGCACTGTTCTGTTGTG | GTGGAAACTCCGTTGAGTGGT |
| Zo1 | CACAGCCTC-CAGAGTTTGACAG | TCCACAGCTGAAGGACTCACAG |
| Ocln | TGGATGACTACAGAGAGGAGAGT | TCCTCTTGATGTGCGATAATTTGC |
| Mlck | AAAAGCCCCATGTGAAACCTTA | TGATTGGTCATCCTTGAACCAG |
| Jama | AGGTCATTTACAGCCAGCCC | AAATGGACAACGGAGGAGCC |
| Muc2 | TGTCCCGACTTCAACCCAAG | TCTGGTTTTGAGGGATGCATGT |
| Reg3g | CCTCCATGATCAAAAGCAGTGG | GGATTCGTCTCCCAGTTGATGT |
| Reg3b | TGGTTTGATGCAGAACTGGC | TGGAGGACAAGAATGAAGCCT |
| Defb1 | ACTCATTACTTTCTCCTGGTGATG | TCCAAGACTTGTGAGAATGCCA |
| Defb2 |  |  |
| Xbp1s | GAACATCTTCCCATGGACTC | CCCAAAAGGATATCAGACTCAG |

**Supplementary Table 3**:

Samples were specifically diluted in the chromatographic H_2_O/CH_3_OH/CH_3_CO_2_H 95/5/0.1 (v/v/v) mobile phase A according to their osmolality measured using a freezing-point osmometer (Loeser Messtechnik, Berlin, Germany). After dilution, samples were centrifuged at 9500g for 10 min. The supernatant was collected and transferred into vials to be directly analyzed by UHPLC-HRMS. A volume of 10 µL was then injected in an ELUTE UHPLC system from Bruker (Bremen, Germany), using H_2_O/CH_3_OH/CH_3_CO_2_H 95/5/0.1 (v/v/v) as mobile phase A and CH3OH/CH3CO2H 100/0.1 (v/v) as mobile phase B, at a flow rate of 0.3 mL/min. The following gradient program was used: from 0 to 30 min: 0% to 100% of B; from 30min to 34 min: 100% B; from 35 to 40 min: 100% A. The separation was achieved at 40°C with a hypersil Gold C18 column (100 × 2.1 mm, 1.9 µm) from Thermo Scientific (Les Ulis, France). The following parameters of electrospray were applied: nebulizer pressure (N2): 3 bar, dry gas flow rate (N2): 10 L·min^−1^, dry temperature 250°C, capillary voltage 3.6 kV for the negative mode and 4.5 kV for the positive mode. High resolution mass spectra were acquired with a TIMS-ToF Flex Q-ToF mass spectrometer from Bruker (Bremen, Germany), between m/z 80 and 800. Samples were analyzed randomly. A QC sample consisted of the pool of all samples of exposed animals was analyzed 7 times and distributed throughout the sequence of injections, to confirm that no signal deviation nor retention time deviation occurred. For metabolite identification, tandem mass spectrometry (MS/MS) experiments were conducted on the QC sample using the PASEF mode at a collision energy ramp between 20 and 50 eV. Ion mobility separation was set up with custom mode from 1/K0 0.45 V·s/cm^2^ to 1.45 V·s/cm^2^ using a ramp time of 100 ms. An internal mass calibration was achieved with every injection, using a mixture of formate clusters (25%) and TuningMix (Agilent, Les Ulis, France) (75%).

Raw data were processed using MetaboScape (version 2024) (Bruker, Bremen, Germany). Boscalid, thiabendazole, cyhalothrin, imazalil, and a list of their phase 1 and phase 2 metabolites were screened into extracted data. Their annotation was based the nomenclature for identification levels from the metabolomics standard initiative (Sumner et al. 2007) according the following parameters: (a) retention time similar as a standard of parent pesticides, (b) *m/z* similar to the theoretical one, (c) isotopic pattern similar to the theoretical one, (d) MS/MS similar to the standard, (e) MS/MS similar to that in the experimental databases (MassBank, HMDB), (f) MS/MS similar to the *in silico* spectrum obtained with MetFrag, (g) annotated in both ionization modes, (h) CCS in accord with the structure calculated by MetaboScape, and (i) not detected in control animals. Annotated metabolites were quantified according to their MS intensities measured by MetaboScape, and only if they were detected in more than 50% of replicates.

| Name | Chemical Formula | Identification level | Annotation parameters | International Chemical Identifier (InChI) |
| --- | --- | --- | --- | --- |
| Boscalid 5-hydroxy glucuronide | C24H20Cl2N2O8 | 2 | b, c ,f, g, h, i | InChI=1S/C24H20Cl2N2O8/c25-12-5-3-11(4-6-12)15-10-13(35-24-19(31)17(29)18(30)20(36-24)23(33)34)7-8-16(15)28-22(32)14-2-1-9-27-21(14)26/h1-10,17-20,24,29-31H,(H,28,32)(H,33,34) |
| Boscalid 5-hydroxy sulfate | C18H12Cl2N2O5S | 2 | b, c, f, i | InChI=1S/C18H12Cl2N2O5S/c19-12-5-3-11(4-6-12)15-10-13(27-28(24,25)26)7-8-16(15)22-18(23)14-2-1-9-21-17(14)20/h1-10H,(H,22,23)(H,24,25,26) |
| Boscalid parent | C18H12Cl2N2O | 1 | a, b, c, d, h, i | InChI=1S/C18H12Cl2N2O/c19-13-9-7-12(8-10-13)14-4-1-2-6-16(14)22-18(23)15-5-3-11-21-17(15)20/h1-11H,(H,22,23) |
| Cyhalothrin 3-[(Z)-2-chloro-3,3,3-trifluoroprop-1-enyl]-2,2-dimethylcyclopropane-1-carboxylic acid glucuronide | C15H18ClF3O8 | 2 | b, c, f, h, i | InChI=1S/C15H18ClF3O8/c1-14(2)4(3-5(16)15(17,18)19)6(14)12(25)27-13-9(22)7(20)8(21)10(26-13)11(23)24/h3-4,6-10,13,20-22H,1-2H3,(H,23,24)/b5-3- |
| Imazalil 1-(2,4-dichlorophenyl)-2-imidazol-1-ylethanol glucuronide | C17H18Cl2N2O7 | 3 | b, c, i | InChI=1S/C17H18Cl2N2O7/c18-8-1-2-9(10(19)5-8)11(6-21-4-3-20-7-21)27-17-12(15(24)25)13(22)14(23)16(26)28-17/h1-5,7,11-14,16-17,22-23,26H,6H2,(H,24,25) |
| Thiabendazole 5-hydroxy glucuronide | C16H15N3O7S | 2 | b, c, f, g, h, i | InChI=1S/C16H15N3O7S/c20-10-11(21)13(15(23)24)26-16(12(10)22)25-6-1-2-7-8(3-6)19-14(18-7)9-4-27-5-17-9/h1-5,10-13,16,20-22H,(H,18,19)(H,23,24) |
| Thiabendazole 5-hydroxy + sulfate | C10H7N3O4S2 | 2 | b, c, f, g, h, i | InChI=1S/C10H7N3O4S2/c14-19(15,16)17-6-1-2-7-8(3-6)13-10(12-7)9-4-18-5-11-9/h1-5H,(H,12,13)(H,14,15,16) |
